# Temporal patterning of the central nervous system by a shared transcription factor code

**DOI:** 10.1101/2020.11.10.376491

**Authors:** Andreas Sagner, Isabel Zhang, Thomas Watson, Jorge Lazaro, Manuela Melchionda, James Briscoe

## Abstract

The molecular mechanisms that ensure the reproducible generation of neuronal diversity in the vertebrate nervous system are incompletely understood. Here we provide evidence of a temporal patterning program consisting of cohorts of transcription factors expressed in neurons generated at successive developmental timepoints. This program acts in parallel to spatial patterning, diversifying neurons throughout the nervous system and in neurons differentiated in-vitro from stem cells. We demonstrate the TGFβ signalling pathway controls the pace of the temporal program. Furthermore, targeted perturbation of components of the temporal program, Nfia and Nfib, reveals their requirement for the generation of late-born neuronal subtypes. Together, our results provide evidence for the existence of a previously unappreciated global temporal program of neuronal subtype identity and suggest that the integration of spatial and temporal patterning programs diversifies and organises neuronal subtypes in the vertebrate nervous system.

## Introduction

In mammals, the function of the nervous system depends on hundreds of molecularly and functionally distinct cell types (Zeng and Sanes, 2017). This diversity requires the generation of different neuronal subtypes at the right place, time and quantity during development, which in turn, guides the wiring of functioning neural circuits. The molecular mechanisms that direct the specification of distinct neuronal classes at characteristic positions, by subdividing the developing nervous system into topographical territories, have received considerable attention (Jessell, 2000; Philippidou and Dasen, 2013). However, even within the same region of the nervous system, most neuronal classes can be further partitioned into distinct subtypes based on molecular and functional properties (Bikoff et al., 2016; Gabitto et al., 2016; Häring et al., 2018; Manno et al., 2020; Sathyamurthy et al., 2018; Zeisel et al., 2018), suggesting that spatial patterning programs are not sufficient to account for the diversity of neuronal subtypes observed in the nervous system.

Temporal mechanisms – the sequential production of different cell types at the same location – have been proposed to contribute to the generation of cell type diversity (Holguera and Desplan, 2018; Kohwi and Doe, 2013). In the Drosophila nervous system, individual neuroblasts produce a characteristic temporal series of distinct neuronal subtypes (Doe, 2017). Similar mechanisms have been documented in some regions of the vertebrate nervous system (Cepko, 2014; Holguera and Desplan, 2018; Oberst et al., 2019). For example, in the cortex distinct subtypes of glutamatergic neurons are sequentially generated (Jabaudon, 2017; Telley et al., 2019), in the hindbrain first motor neurons (MNs) and later serotonergic neurons are generated from the same set of progenitors (Pattyn et al., 2003), while in the midbrain, the production of ocular MNs is followed by red nucleus neurons (Deng et al., 2011). Moreover, progenitors throughout the nervous system typically produce neurons first and later generate glial cells such as astrocytes and oligodendrocytes (Miller and Gauthier, 2007; Rowitch and Kriegstein, 2010). However, whether temporal programs are a universal feature of neuronal subtype specification in the vertebrate nervous system and whether these are implemented by common or location specific mechanisms is unclear.

The vertebrate spinal cord is an experimentally tractable system to address the basis of neuronal diversity. In this region of the nervous system, neurons process sensory inputs from the periphery relaying the information to the brain or to motor circuits that control and coordinate muscle activity. The temporally stratified generation of some of these neuronal subtypes has been documented, including inhibitory and excitatory neurons located in the dorsal horn as well as ventral motor and interneurons (Benito-Gonzalez and Alvarez, 2012; Deska-Gauthier et al., 2020; Hayashi et al., 2018; Hollyday and Hamburger, 1977; Luxenhofer et al., 2014; Müller et al., 2002; Sockanathan and Jessell, 1998; Stam et al., 2012). Furthermore, the birth order of neurons seems to control the specificity of neuronal connectivity, with flexor and extensor muscle premotor interneurons born at different timepoints during development (Tripodi et al., 2011). Nevertheless, a comprehensive picture is lacking and the genetic programs that orchestrate the temporal patterning of the spinal cord are largely unclear. To this end, we recently characterized the emergence of neuronal diversity in the embryonic spinal cord (Delile et al., 2019). This revealed cohorts of transcription factors (TFs) that further partition all major neuronal subtypes. Moreover, the onset of expression of these different cohorts occurs at characteristic timepoints during the neurogenic period of spinal cord development (Figure 1A). In all domains, the earliest neurons express Onecut-family TFs, intermediate neurons express Pou2f2 and Zfhx2-4, while late-born neurons express Nfia/b/x and Neurod2/6 (Delile et al., 2019; Sagner and Briscoe, 2019). This suggests the existence of a previously unappreciated temporal dimension to neuronal subtype generation in the spinal cord.

**Figure 1:**
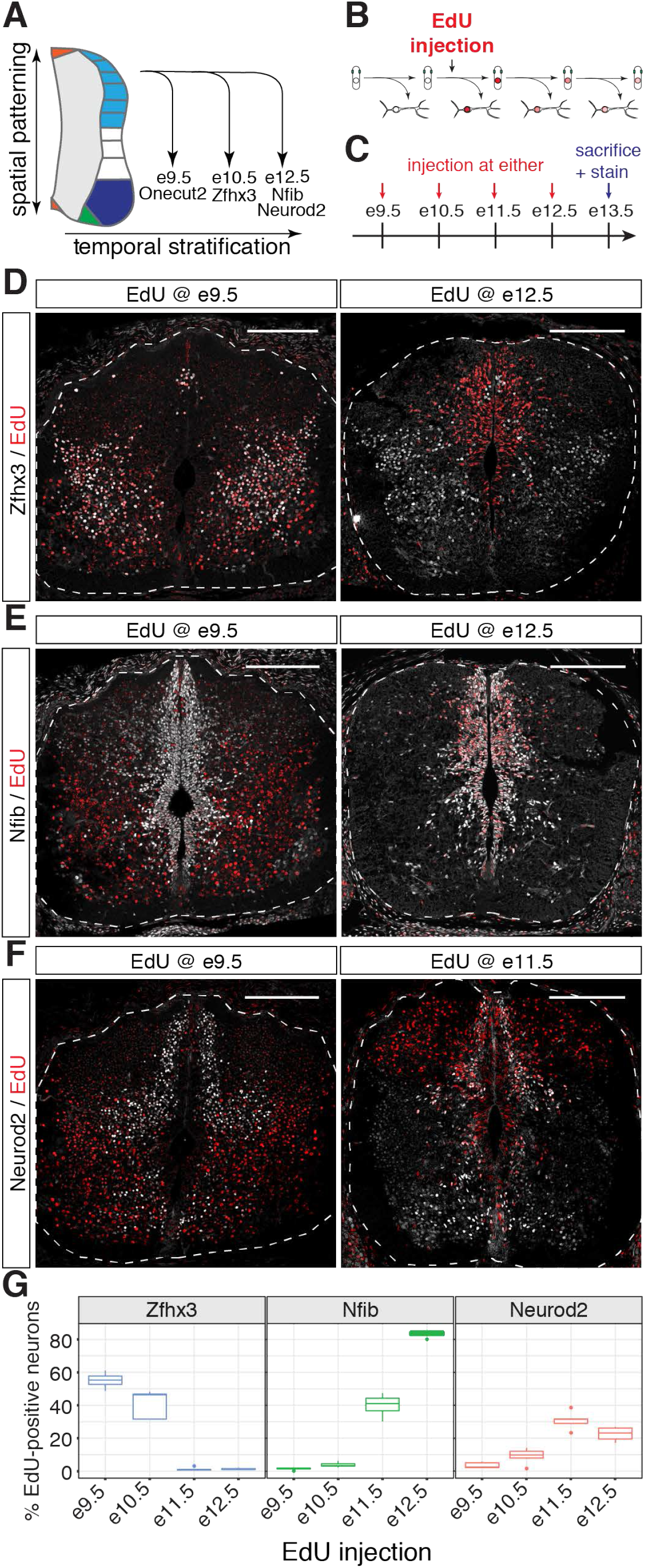
Distinct birth-dates of neurons expressing different temporal TFs (see also Figures S1 and S2) (A) Distinct cohorts of TFs are induced at different developmental stages in neurons from all dorsal-ventral domains in the spinal cord (B) Scheme depicting EdU-birthdating of neurons. (C) Dams were injected with EdU at e9.5, e10.5, e11.5 or e12.5 and embryos collected at e13.5. Colocalization between EdU and temporal TFs was then assessed in spinal cord cryosections. (D) Zfhx3-positive neurons are labelled by EdU, when EdU is administered at e9.5, but not at e12.5. (E) EdU labels Nfib-positive neurons when administered at e12.5, but not at e9.5. (F) Neurod2-postive neurons are labelled when EdU is administered at e11.5, but not when EdU is administered at e9.5. (G) Percentage of EdU-positive neurons labelled by Zfhx3, Nfib and Neurod2 in the spinal cord. Scale bars in D,E,F = 200 µm

Although the role of these TFs had not been conceptualized as part of a globally coordinated temporal code, some have been shown to specify subpopulations of neurons in individual domains in the spinal cord. Onecut TFs, for example, are required in early-born V1 and MNs for the specification of Renshaw cells and medial lateral motor column (LMCm) neurons respectively (Roy et al., 2012; Stam et al., 2012). Furthermore, Onecut TFs and Pou2f2 control the distribution of neurons from multiple dorsal-ventral domains (Harris et al., 2019; Kabayiza et al., 2017; Masgutova et al., 2019). Neurod2/6 control neuropeptide expression in inhibitory neurons in the dorsal horns of the spinal cord (Bröhl et al., 2008) and characterization of V2a neuron heterogeneity revealed that Zfhx3 and Neurod2/Nfib divide this neuronal class into a lateral and medial population (Hayashi et al., 2018). Similar to the spinal cord, Onecut, Pou2f2 and Nfi-TFs label early and late born neuronal subtypes in the retina and are required for their generation (Clark et al., 2019; Javed et al., 2020; Sapkota et al., 2014) and Pou2f2, Zfhx3 and Nfi TFs define distinct subpopulations of Pitx3-positive neurons born from the midbrain floor plate including dopaminergic neurons (Tiklová et al., 2019). These observations raise the possibility that this temporal TF code is conserved in large parts of the central nervous system.

TGFβ signalling has been implicated in the timing of developmental temporal switches in the nervous system (Dias et al., 2014; Rossi and Desplan, 2020). The transition from MN to serotonergic neurons and from ocular MNs to red nucleus neurons is accelerated by TGFβ signalling (Dias et al., 2014). TGFβ signalling also promotes the expression of the late progenitor marker Nfia in neurogenic neural stem cells (Tchieu et al., 2019). Furthermore, Growth differentiation factor 11 (Gdf11), a ligand of the TGFβ family that signals via Activin receptors (Andersson et al., 2006; Paul Oh et al., 2002), has been implicated in the timing of MN subtype generation and onset of gliogenesis in the spinal cord (Shi and Liu, 2011). TGFβ signalling is also important for controlling the timing of fate switches in the Drosophila nervous system (Rossi and Desplan, 2020), raising the possibility that it may serve as a general timer for the sequential generation of cellular subtypes.

Here, we demonstrate by EdU-birthdating that a set of TFs comprise a temporal TF code that identifies neurons based on their timepoint of birth. We find that the same sequence of TF expression applies throughout the brain and for stem cell derived in-vitro generated neurons with defined dorsal-ventral and axial identities. We also document a temporal patterning code for progenitors throughout the nervous system and provide evidence that TGFβ signalling controls the pace of the temporal program. Finally, to characterize the genetic programs that control the temporal specification of neurons, we perturb the function of the TFs Nfia and Nfib and show that their activity is required for the generation of late neuronal subtypes. Taken together, our data reveal conserved temporal patterning of neurons and progenitors in large parts of the nervous system that is under the control of the TGFβ signalling pathway and suggest a close link between the developmental programs that control the switch from neuro- to gliogenesis and the specification of neuronal diversity.

## Results

### EdU-birthdating reveals a temporal TF code in spinal cord neurons

We previously identified cohorts of TFs that are expressed in multiple subsets of neurons in the spinal cord. As the onset of expression of these TFs occurred at different times during development, we speculated that they subdivide neurons in the spinal cord based on their timepoint of birth (Delile et al., 2019) (Figure 1A). We and others have demonstrated before that Onecut TFs are expressed in early-born neurons and that their expression is rapidly extinguished as neurons mature (Delile et al., 2019; Kabayiza et al., 2017; Luxenhofer et al., 2014; Roy et al., 2012; Stam et al., 2012). We therefore focused on the TFs Zfhx3, Nfib and Neurod2, which start to be expressed in neurons at intermediate or late stages during the neurogenic period respectively, and analysed the birth date of neurons expressing these TFs by EdU incorporation (Figure 1B). Pregnant dams were injected with EdU at embryonic day (e)9.5, e10.5, e11.5 or e12.5 (Figure 1C). Embryos were collected at e13.5 and forelimb-level spinal cord cryo-sections assayed for colocalization between EdU and Zfhx3, Nfib and Neurod2 in neurons (Figure 1D-F and Figure S1).

Consistent with the hypothesis of a temporal TF code, a high proportion of EdU-labelled neurons expressed Zfhx3, when EdU was administered at e9.5 and e10.5, while there was little if any colocalization between EdU and Zfhx3 when EdU was given at later timepoints (Figures 1G and S1A). By contrast, the proportion of EdU+ neurons expressing Nfib continually increased. Few EdU-positive neurons expressed Nfib when EdU was administered before e11.5, but more than 80% of EdU+ neurons were positive for Nfib when it was given at e12.5 (Figures 1G and S1B). Neurod2 followed a similar trend to Nfib until e11.5 (Figures 1G and S1C), consistent with the high degree of co-expression between these genes (Delile et al., 2019). However, the proportion of Neurod2-positive neurons decreased when EdU was given at e12.5 (Figures 1G and S1C). This may be due to the relatively late onset of Neurod2 expression after neuronal differentiation. Furthermore, Neurod2 is not expressed in late-born dorsal excitatory neurons (Bröhl et al., 2008), which are generated at high frequency during late neurogenic stages in the spinal cord (Wildner et al., 2006).

The mutually exclusive birth dates of Zfhx3 and Nfib/Neurod2-positive neurons suggest that these TFs label largely non-overlapping subsets of neurons. To test this prediction directly, we stained e13.5 spinal cord sections for either Zfhx3 and Nfib or Zfhx3 and Neurod2 (Figure S2). Although each of these markers labelled a large number of neurons, the expression of Zfhx3 and Nfib or Zfhx3 and Neurod2 was mutually exclusive. Taken together, the birth dates of neurons expressing different TFs closely matches our previous description of the onset of expression of these TFs from scRNAseq data (Delile et al., 2019). These results are consistent with a model in which Zfhx3 is specifically expressed and maintained in neurons born before e11.5 but not in later-born neurons, which instead express Neurod2/6 and Nfi-family TFs. These results further argue against sequential expression of these TFs during neuronal maturation because in such a model TFs with early onset of expression would be specific for early maturation stages and would thus, contrary to our observations, be expected to be labelled by EdU given at late developmental timepoints. We therefore conclude that these data provide clear evidence that these TFs comprise a temporal code and label distinct subsets of neurons based on their timepoint of birth in the spinal cord.

### Conservation of the temporal TF code in other regions of the nervous system

Similar to the spinal cord, Pou2f2 and TFs of the Onecut and Nfi-families are required for the generation of early and late-born neurons in the retina (Clark et al., 2019; Javed et al., 2020; Sapkota et al., 2014). We therefore speculated that the temporal TF code is preserved in the retina. To test this hypothesis, we analysed a published scRNAseq time course of mouse retina development (Clark et al., 2019) (Figure S3). Performing dimensionality reduction by Uniform Manifold Approximation and Projection (UMAP) on the data from pre- and perinatal stages (e14, e16, e18, P0) resulted in clear trajectories from retinal progenitors to horizontal cells, amacrine cells, retinal ganglion cells and cone and rod photoreceptors (Figure S3A,B). Examining Onecut2, Pou2f2, Zfhx3 and Nfib revealed different expression of these genes along these differentiation trajectories (Figure S3C). As expected, Onecut2 was strongly enriched in horizontal cells, an early born cell type in the retina, although some expression was also observed in retinal ganglion cells, amacrine cells, and cones. Nfib expression was largely restricted to late progenitors and rods (Figure S3C). By contrast, Pou2f2 and Zfhx3 were enriched in amacrine and retinal ganglion cells. Furthermore, both genes were expressed in subsets of retinal progenitors.

To further characterize the expression of Onecut2, Pou2f2, Zfhx3 and Nfib genes in retinal neurons, we plotted their expression levels in the individual classes of neurons stratified by developmental stage (Figure S3D). This analysis revealed a clear link between the expression of these TFs and developmental stage. Onecut2 was enriched in amacrine cells, retinal ganglion cells and cones at e14 (Figure S3D). Zfhx3 was absent at this stage but was enriched in amacrine and retinal ganglion cells at e18 and P0 (Figure S3D). These data support the hypothesis that the temporal TFs are expressed in different retinal cell types born at distinct timepoints and raise the possibility that the expression of these genes further subdivide distinct classes of retinal neurons based on their birth dates.

Nfi TFs have been previously shown to be expressed in neurons in the forebrain including the cortex, thalamus and hippocampus (Piper et al., 2010; Plachez et al., 2008), while Zfhx3 has been implicated in controlling circadian function of the suprachiasmatic nucleus (Parsons et al., 2015). Moreover, Pou2f2, Zfhx3 and Nfi TFs are expressed in subpopulations of neurons born from the midbrain floorplate (Tiklová et al., 2019). These results raised the possibility that the sequential expression of the temporal TFs might be broadly preserved throughout the developing nervous system. To test this, we first turned our attention to available scRNAseq timecourse data from the developing forebrain, midbrain and hindbrain (Manno et al., 2020). Plotting the dynamics of Onecut1-3, Pou2f2, Zfhx3/4, Nfia/b and Neurod2/6 in neurons between e8.5 and e14 revealed a striking conservation of the expression dynamics of these TFs. Expression of Onecut-family TFs preceded Pou2f2 and Zfhx3/4, while Nfia/b and Neurod2/6 were only expressed at high levels at later stages (Figure 2A). To experimentally validate these predictions, we turned to immunofluorescent analysis of Onecut2, Pou2f2, Zfhx3 and Nfib in hind- and midbrain cryo-sections from different developmental stages (Figure 2B-H). In both tissues, the majority of neurons expressed Onecut2 but not Pou2f2 at early developmental stages (e9.5 or e10.5 respectively) and both genes were expressed in largely non-overlapping populations of neurons one day later (Figure 2B,E,F). Furthermore, in both tissues a large proportion of neurons expressed Zfhx3 at e11.5, while Nfib expression was confined to neural progenitors at this stage (Figure 2C,G,H). At e13.5 Nfib-positive cells, which had lost the expression of the progenitor marker Sox2, were detected in the mantle layer of both tissues where postmitotic neurons reside (Figure 2C,G,H). To test if these Nfib-expressing cells are neurons, we co-stained hindbrain sections for Phox2b, which is expressed in different populations of hindbrain neurons (Dubreuil et al., 2009). This analysis revealed colocalization between Phox2b/Zfhx3 and Phox2b/Nfib in individual nuclei (Figure 2D), suggesting that Nfib indeed labels late-born neurons in the hindbrain. These results suggest the existence of a conserved temporal patterning program that subdivides neurons based on their timepoint of birth throughout the developing nervous system.

**Figure 2:**
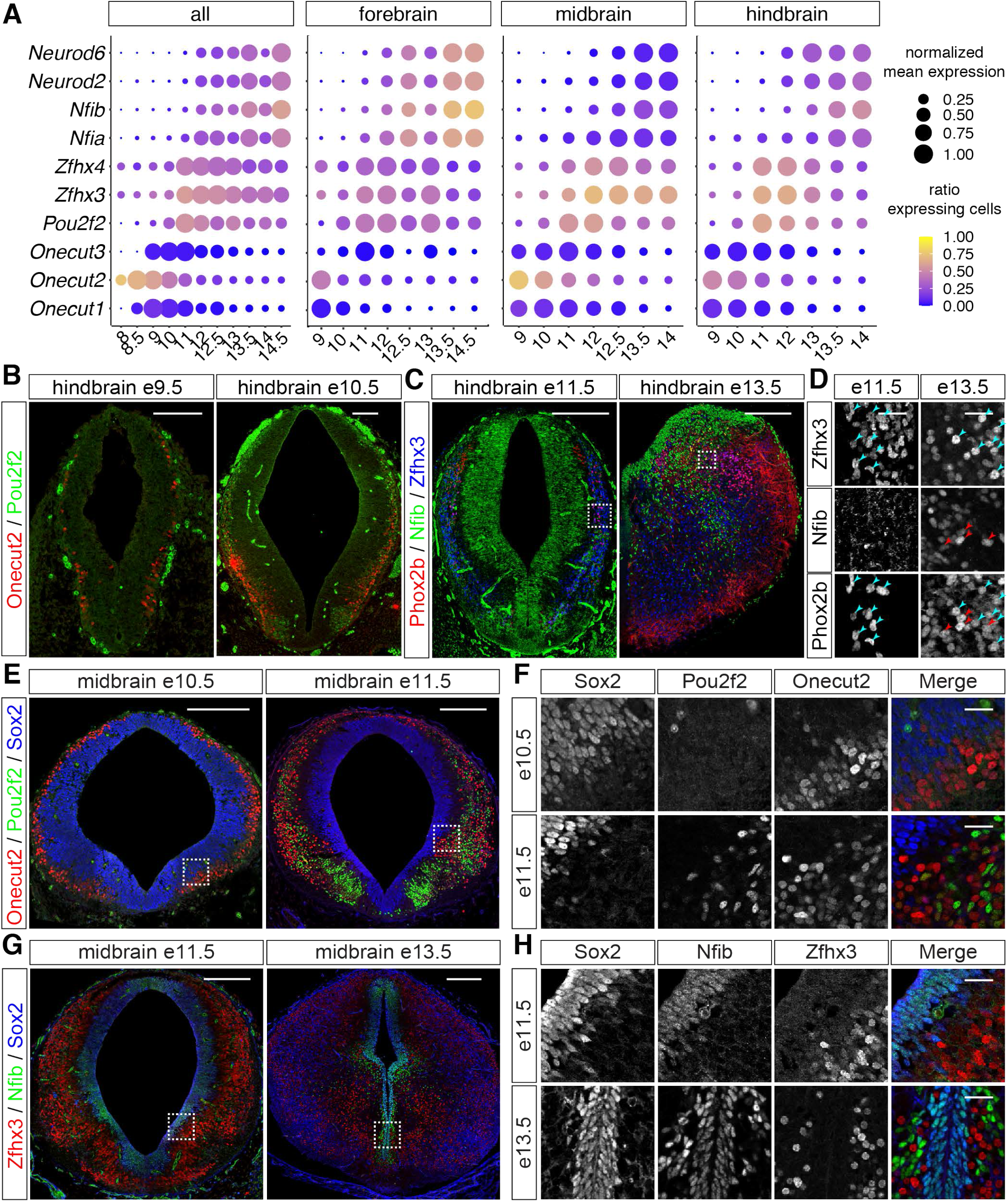
The temporal TF code is conserved at different rostral-caudal levels of the nervous system (see also Figure S3) (A) Expression of temporal TFs in scRNAseq data (Manno et al., 2020) from the developing forebrain, midbrain and hindbrain suggests conservation of temporal patterning in these parts of the nervous system. (B-D) Conservation of temporal patterning in the hindbrain. (B) Onecut2, but not Pou2f2 is expressed in hindbrain neurons at e9.5, while both TFs label distinct populations of neurons at e10.5. (C) Zfhx3, but not Nfib labels neurons at e11.5. These TFs label distinct populations of neurons at e13.5. (D) Zfhx3 and Nfib label distinct subsets of Phox2b-positive neurons in the hindbrain at e13.5. (E-H) Conservation of temporal patterning in the midbrain. (F,H) show higher magnification images of the regions outlined in (E,G) respectively. (E,F) Onecut2, but not Pou2f2, labels neurons in the midbrain at e10.5. Both TFs label distinct subsets of neurons at e11.5. (G,H) Zfhx3 labels neurons at e11.5, while Nfib expression is restricted to neural progenitors. At e13.5 Zfhx3 and Nfib label distinct subsets of neurons in the midbrain at e13.5. Scale bars = 100 µm (B), 200 µm (C,E,G) or 25 µm (D, F, H)

### Dopaminergic neurons are a temporal neuronal subtype generated in the midbrain

We next investigated if this temporal TF code is responsible for the establishment of neuronal populations with specific functions. Dopaminergic neurons are a neuronal population of medical interest because their degeneration causes Parkinson’s disease. During development these neurons are born from the midbrain floor plate and can be discriminated based on the expression of the TFs Lmx1a, Lmx1b and Pitx3 as well as the enzymes Tyrosine hydroxylase (TH) and the dopamine transporter Slc6a3 (also known as Dat). Strikingly, previous characterization of neurons generated from the midbrain floor plate using a Pitx3 transgenic reporter suggested that these neurons can be broadly subdivided into Nfia/b and Zfhx3 expressing subsets. The Zfhx3-positive population expresses dopaminergic neuron markers such as Slc6a3 and high levels of TH (Tiklová et al., 2019). By contrast, the Nfi-positive population lacked the molecular machinery for the synthesis of dopamine and expressed markers characteristic for excitatory neurons such as Slc17a6 (also known as vGlut2) (Tiklová et al., 2019). These findings, in combination with our observation that Zfhx3 and Nfi TFs define temporal neuronal populations in the midbrain suggest that midbrain dopaminergic neurons may constitute a temporal neuronal subtype born from the midbrain floor plate.

We therefore examined if Zfhx3-positive neurons are generated before Nfia/b positive neurons from the midbrain floor plate. Assays at e11.5 revealed widespread expression of Zfhx3 in floor plate-derived Lmx1b-positive neurons (Figure 3A). At this stage Nfib expression just commenced in Sox2+ neural progenitors (Figure 3B). In contrast, at e13.5 numerous Nfib-positive neurons expressing Lmx1b were found in the vicinity of the midbrain floor plate (Figure 3C,D), likely corresponding to the *N-Dat*^*low*^ population (Tiklová et al., 2019). Zfhx3-positive neurons at this stage had migrated to a more lateral position (Figure 3C). These neurons co-expressed the Zfhx TFs, Zfhx3 and Zfhx4, and also increased levels of TH (Figure 3E,F), suggesting these populations correspond to the *AT-Dat*^*high*^, *T-Dat*^*high*^ and *VT-Dat*^*high*^ neurons described by Tiklová et al., 2019. These conclusions are also consistent with previous birth-dating experiments that concluded that the majority of TH-positive dopaminergic neurons are born before and around e12.5 (Bayer et al., 1995; Bye et al., 2012). Taken together, these data suggest that the sequence of temporal TF expression is preserved in neurons derived from the midbrain floor plate, that the expression of different temporal TFs correlates with the acquisition of different neuronal subtype identities in these neurons and that dopaminergic neurons correspond to the Zfhx3-positive temporal neuronal population.

**Figure 3:**
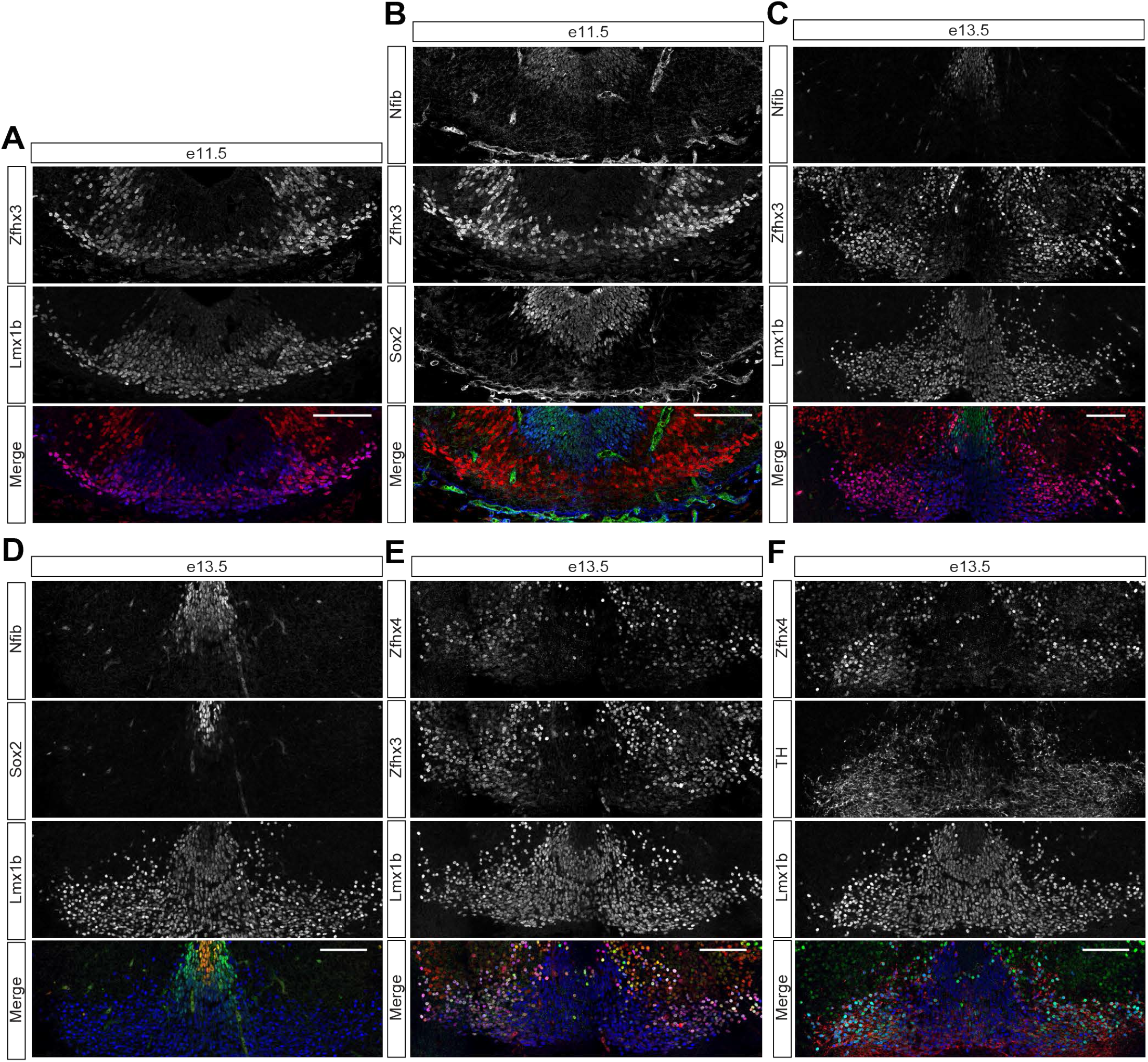
Midbrain dopaminergic neurons are a temporal population of neurons derived from the midbrain floor plate. (A) Coexpression of Zfhx3 and Lmx1b in neurons derived from the midbrain floor plate at e11.5 (B) Nfib is restricted to Sox2+ neural progenitors in the ventral midbrain at e11.5 (C) Mutually exclusive expression of Zfhx3 and Nfib in Lmx1b-positive neurons at e13.5 (D) Nfib labels Lmx1b-positive neurons directly adjacent to Sox2-positive progenitors at e13.5 (E) Colocalization between Zfhx3 and Zfhx4 in Lmx1b-positive neurons at e13.5 (F) Zfhx4 labels Lmx1b-positive neurons expressing high-levels of TH at e13.5 Scale bars = 100 µm

### The temporal TF code applies to in-vitro generated midbrain, hindbrain and spinal cord neurons

We next sought to investigate whether the temporal code was preserved in-vitro during the directed differentiation of ES cells to neurons with specific axial and dorsal-ventral identities (Gouti et al., 2014; Metzis et al., 2018; Sagner et al., 2018). We reasoned that, in-vitro putative global signalling cues, originating from distant signalling centres, should be absent.

We examined if the same sequence of temporal TF factor expression can be observed in stem-cell derived neurons with mid- and hindbrain and spinal cord identities. ES cells were differentiated to appropriate identities using established protocols (Gouti et al., 2014) (Figure 4A), as confirmed by real-time quantitative polymerase chain reaction (RT-qPCR) for *Foxg1, Otx2, Hoxa4, Hoxb9* and *Hoxc8* (Figure S4A). As expected, cells differentiated to midbrain identity induced *Otx2*, but not the forebrain marker *Foxg1* or the hindbrain marker *Hoxa4*, which was induced in hindbrain conditions. By contrast, the posterior Hox genes *Hoxb9* and *Hoxc8* were only induced when cells were differentiated to a spinal cord identity. We next assayed the expression of the temporal TFs Onecut2, Zfhx3, Nfia and Neurod2 under these differentiation conditions by flow cytometry from days 6 to 13 (Figures 4B and S4B). The overall expression dynamics of these markers observed in-vivo were preserved under the different conditions. Most neurons expressed Onecut2 at days 6 and 7, while the proportion of Zfhx3-positive neurons increased between days 7 and 9 and Nfia and Neurod2-positive neurons were typically not detected before day 11. These results closely resemble our previous observations of the temporal patterning of neurons in the developing nervous system.

**Figure 4:**
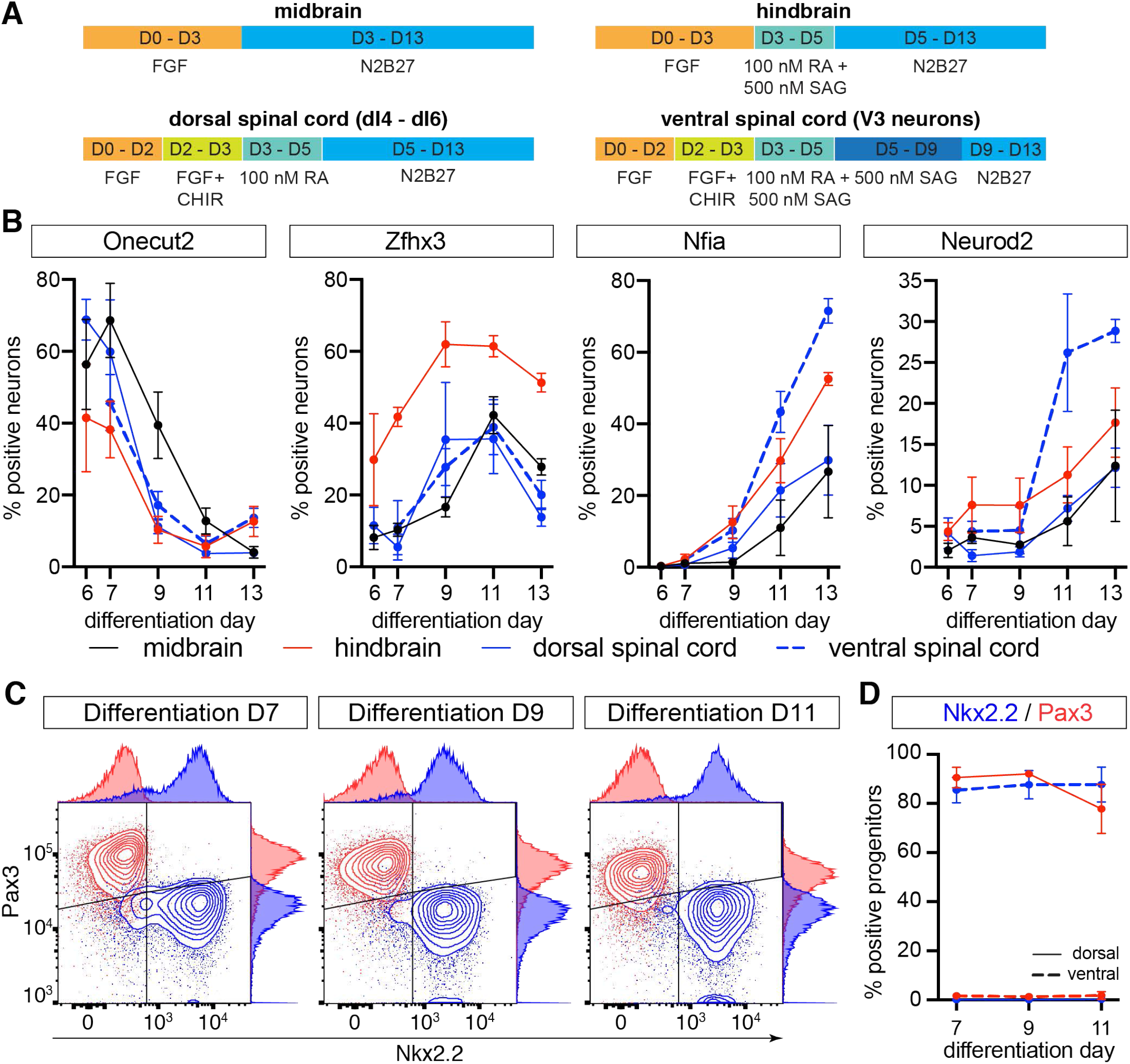
Conservation of the temporal TF code in stem-cell derived neurons with different axial and dorsal-ventral identities (see also Figure S4) (A) Schematics of the differentiation protocols for the generation of progenitors and neurons with different axial and dorsal-ventral identities (B) Flow cytometry analysis of temporal TF expression indicates that neurons with different axial and dorsal-ventral identities display the same temporal progression in-vitro as in-vivo. (C) Flow cytometry analysis of Nkx2.2 and Pax3 expression in neural progenitors in dorsal and ventral spinal cord differentiations. (D) Percentage of neural progenitors expressing Pax3 and Nkx2.2 in ventral and dorsal spinal cord differentiations between days 7-11.

We next investigated if the progression of the temporal TF code is preserved in neurons with different dorsal-ventral identities. We have previously demonstrated that exposure of spinal cord progenitors to appropriate concentrations of the Sonic Hedgehog (Shh) pathway agonist (SAG) promotes the generation of progenitors and neurons with different dorsal-ventral identities (Sagner et al., 2018). We therefore focused on the spinal cord condition and either ventralized cells by exposing them from day 3 to day 9 to 500 nM SAG, or dorsalised them in the absence of SAG. Samples for flow cytometry were collected at days 7, 9, and 11 (Figure 4A). Consistent with our previous observations (Sagner et al., 2018), in the absence of Shh pathway activation most progenitors expressed the dorsal progenitor marker Pax3, while prolonged high-level Shh pathway activation leads to the majority of progenitors acquiring a Nkx2.2-positive ventral p3 identity (Figure 4C,D). Consistent with this, most neurons generated in the absence of Shh pathway activation expressed the intermediate dorsal marker Lbx1, while Shh pathway activation lead to the generation of Sim1-positive V3 neurons (Figure S4C). We therefore refer to these conditions as dorsal and ventral, respectively. Assaying the expression of the temporal TFs in neurons in the ventral differentiation condition revealed similar expression dynamics for these markers as previously observed under dorsal spinal cord conditions, although notably a higher proportion of neurons expressed Nfia and Neurod2 at later stages of the differentiations (Figure 4B).

Based on these oberservations, we conclude that the temporal TF code is preserved in in-vitro generated neurons with different axial (midbrain, hindbrain and spinal cord) and dorsal-ventral identities. Furthermore, the time scale over which the temporal patterning unfolds is similar in-vivo and in-vitro, corresponding in both cases to approximately 4-5 days (in-vivo ∼e9.5 – e13.5; in-vitro ∼day 7 – day 11). These results argue against a model in which global signalling cues orchestrate the temporal patterning program. We note however, that this analysis also uncovered reproducible differences in the proportions of neurons expressing the respective markers between the different axial identities. Cells differentiated under hindbrain conditions induced late temporal TFs at a faster pace, while cells under midbrain conditions seemed to progress slowest to a later temporal identity. These differences may be indicative of cell-intrinsic programs that allow progenitors and/or neurons to progress through the temporal TF code at a speed characteristic for their axial identity.

### Conserved temporal patterning of midbrain, hindbrain and spinal cord neural progenitors

The generation of different neuronal subtypes from the same progenitors is arguably best understood in Drosophila. Here, aging neuroblasts sequentially express a series of TFs that define temporal identity windows for the generation of specific neuronal progeny (Doe, 2017). Similar processes are believed to underlie the temporal patterning of tissues in the vertebrate nervous system, however, the transcriptional programs that mediate this process are still relatively poorly understood (Holguera and Desplan, 2018; Oberst et al., 2019). We therefore asked if similar principles apply to the spinal cord. To this end, we analysed our in-vivo scRNAseq data (Delile et al., 2019) to identify TFs that are consistently up- or downregulated in most progenitor domains during the neurogenic period (see Experimental Procedures). This analysis recovered in total 542 genes including 33 TFs (Figure 5A). Inspection of the expression dynamics of these TFs confirmed their differential temporal expression in progenitors from most dorsal-ventral domains (Figure S5A). As expected, this analysis recovered the gliogenic TFs Sox9 and the Nfi TFs (Nfia/b/x) that have previously been shown to be dynamically expressed during this time window in the developing spinal cord (Deneen et al., 2006; Stolt et al., 2003). To address if these transcriptional changes are preserved in progenitors in other regions of the nervous system, we characterized the expression dynamics of the 33 TFs in scRNAseq from the developing fore-, mid- and hindbrain (Manno et al., 2020). This analysis revealed largely preserved expression dynamics of the 33 TFs in these tissues (Figure 5B). We next tested if these genes display the same expression dynamics in neural progenitors with different axial identities in our in-vitro differentiations. Analysis of the gene expression dynamics of the 542 genes and 33 TFs using RNAseq data from in-vitro generated ventral neural progenitors from days 3-10 (Rayon et al., 2020) revealed that the general temporal pattern of gene expression is preserved under these culture conditions (Figure 5C). To test if the same dynamics can be also observed in in-vitro generated progenitors with midbrain, hindbrain or dorsal spinal cord identities, we performed RT-qPCRs for *Lin28a, Lin28b, Nr6a1, Sox9, Npas3, Zbtb20, Nfia, Nfib* and *Hopx* and quantified the proportion of Nfia-positive progenitors by flow cytometry (Figures 5D,E and S5B). These results confirmed shared expression dynamics for these marker genes in in-vitro differentiated neural progenitors with different axial identities. We conclude that, similar to neurons, neural progenitors throughout the nervous system undergo a shared temporal patterning program (Figure 5F).

**Figure 5:**
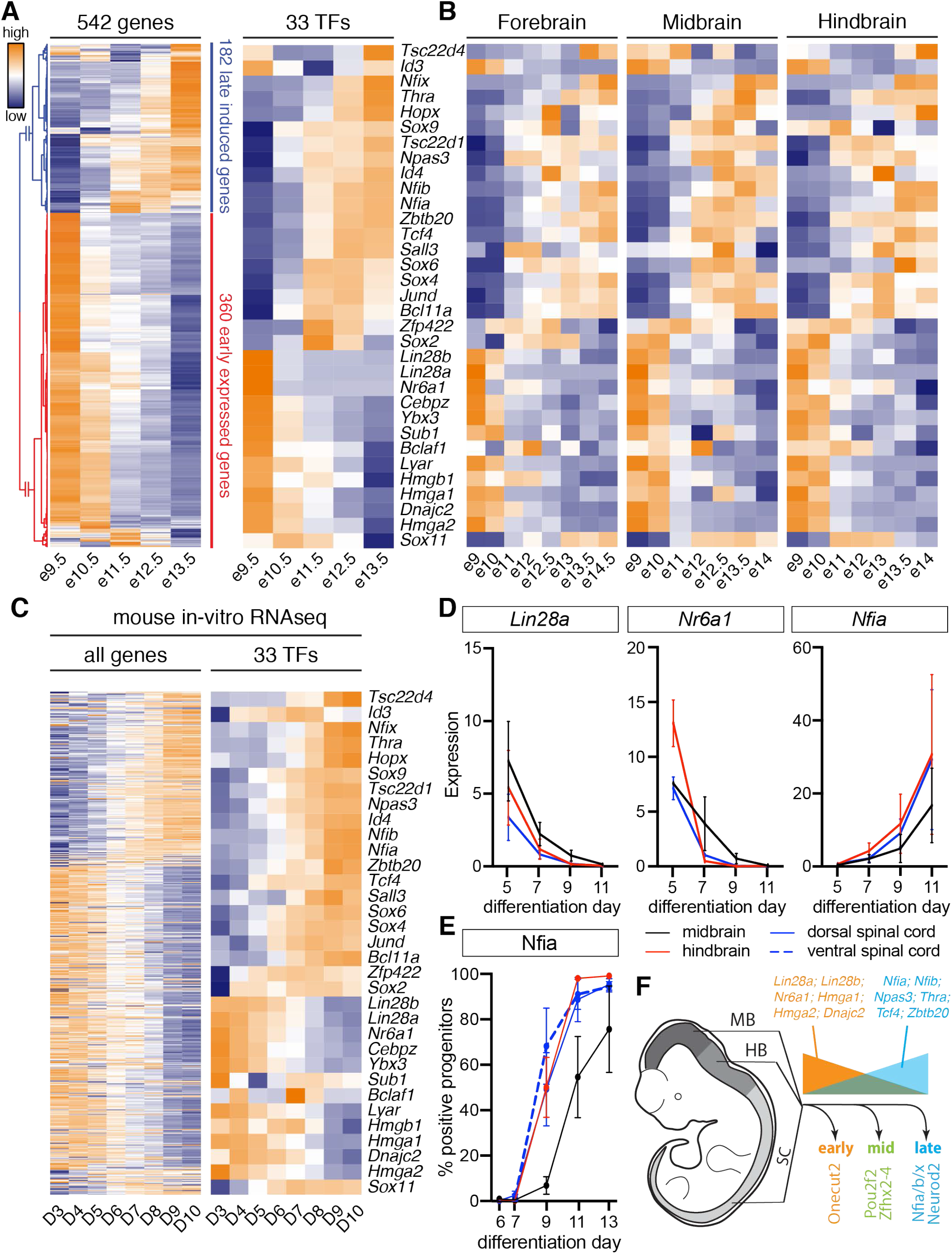
Conserved temporal patterning of neural progenitors throughout the developing central nervous system (see also Figure S5) (A) Differential gene expression analysis using scRNAseq from spinal cord neural progenitors (Delile et al., 2019) identifies 542 genes (left) including 33 TFs (right) that are differentially expressed during the neurogenic period. Heatmap shows log-scaled and z-scored gene expression values for each gene. (B) Characterization of the expression dynamics of the same 33 TFs in scRNAseq from the developing forebrain, midbrain and hindbrain (Manno et al., 2020). (C) Expression dynamics of the 542 genes (left) and 33 TFs (right) in RNAseq data from ventral spinal cord differentiations (Rayon et al., 2020). Heatmap shows log-scaled and z-scored gene expression values for each gene. Order of the genes in both heatmaps is the same as in (A). (D) RT-qPCR analysis of *Lin28a, Nr6a1* and *Nfia* from days 5-11 in in-vitro differentiations with different axial identities reveals conserved expression dynamics of these markers in the in-vitro differentiations. See Figure S5B for quantification of further markers. (E) Quantification of Nfia induction in in-vitro generated neural progenitors with different axial identities by flow cytometry. (F) Conserved temporal patterning of neural progenitors throughout the developing nervous system. Early neural progenitors express markers such as *Lin28a, Lin28b, Nr6a1, Hmga1, Hmga2* and *Dnajc2* (orange), while late progenitors are characterized by the expression of *Nfia, Nfib, Npas3, Thra, Tcf4* and *Zbtb20* (light blue).

### TGFβ controls the pace of the temporal program

The TGFβ signalling pathway controls the timing of the switch from motor neuron to serotonergic neuron production in p3 progenitors in the vertebrate hindbrain and promotes Nfia expression and the formation of glia in neural stem cells (Dias et al., 2014; Tchieu et al., 2019). In the spinal cord, the signalling pathway is active in progenitors during the neurogenic period and several members of the TGFβ family are expressed at early developmental stages in the adjacent notochord, floor plate, mesoderm and at later developmental stages by different populations of neurons (Dutta et al., 2018; Garcia-Campmany and Marti, 2007; Shi and Liu, 2011). We therefore asked if the pathway is active in progenitors in our in-vitro differentiations. To do so, we exposed dorsal neural progenitors from day 5 to the TGFβ signalling inhibitor SB431542 (Inman et al., 2002) and assayed the expression of the target gene *Smad7* 48 and 96 hours later (Garcia-Campmany and Marti, 2007). As expected, pathway inhibition resulted in a significant reduction of *Smad7* expression (Figure 6B). These results confirm that the TGFβ pathway is active in neural progenitors in-vitro and suggest that TGFβ signalling is a good candidate to control the maturation of progenitors and the timing of temporal TF expression in in-vitro generated spinal cord neurons

**Figure 6:**
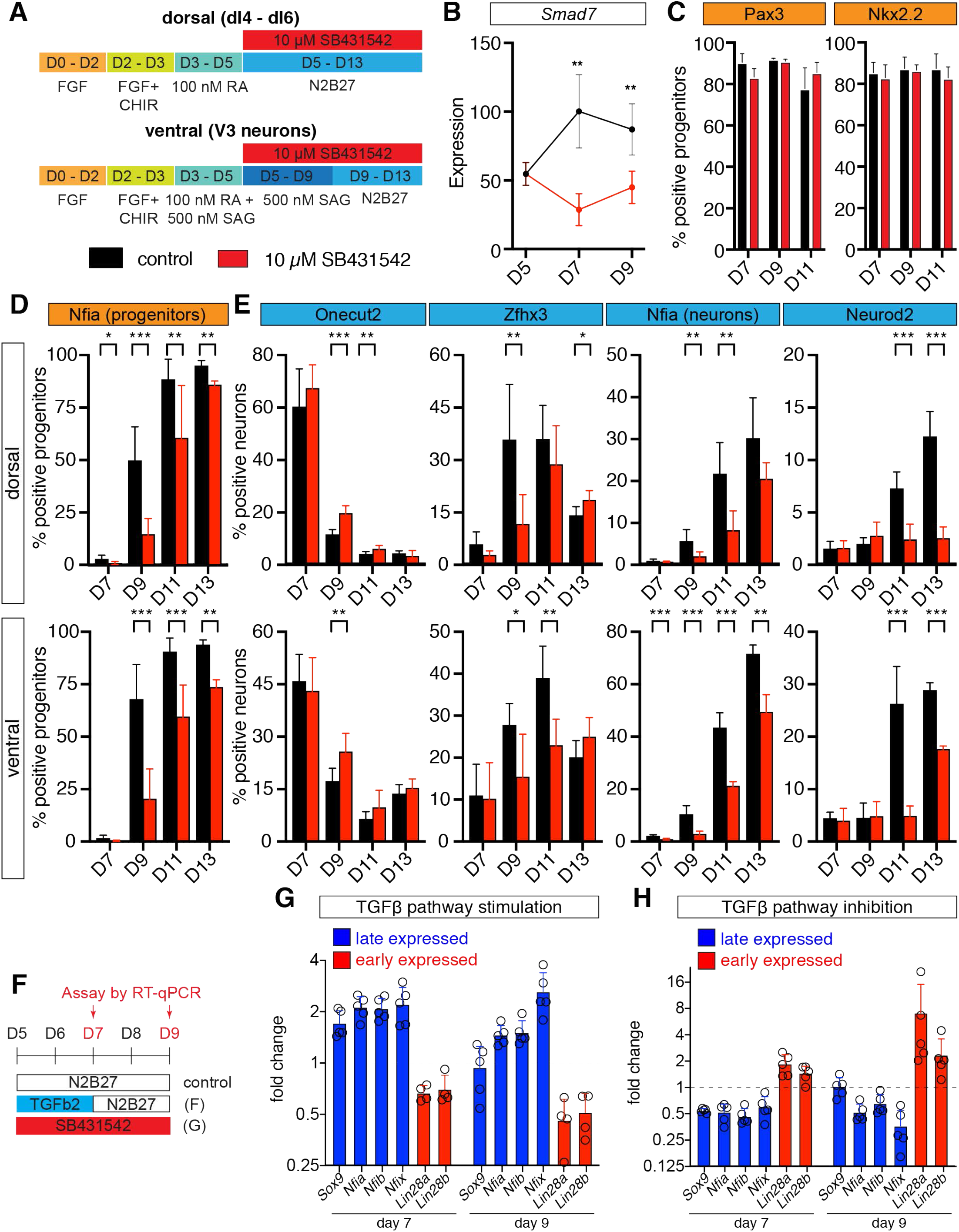
TGFβ signalling influences the timing of temporal TF expression in neurons and progenitors. (A) Schematics of the differentiation protocols for TGFβ pathway inhibition in dorsal and ventral spinal cord conditions (B) Inhibition of TGFβ signalling in dorsal spinal cord conditions causes down-regulation of the TGFβ pathway target gene *Smad7*. (C) TGFβ pathway inhibition does not alter the proportion of progenitors expressing Pax3 in dorsal (left) or Nkx2.2 in ventral (right) conditions. (D) Inhibition of TGFβ signalling delays the induction of Nfia in dorsal and ventral spinal cord neural progenitors. (E) Percentage of neurons expressing the different temporal TFs in the presence and absence of TGFβ pathway inhibition. TGFβ pathway inhibition causes a delay in the induction of the late neuronal markers Zfhx3, Nfia and Neurod2 in neurons. (F) Scheme outlining the differentiation protocol to assess the role of TGFβ pathway activation and inhibition on the temporal patterning of neural progenitors. (G) TGFβ pathway activation causes an earlier induction of the late markers *Sox9, Nfia, Nfib* and *Nfix* and earlier downregulation of *Lin28a* and *Lin28b* by RT-qPCR. (H) TGFβ pathway inhibition has the opposite effect on the expression of these markers.

To test this hypothesis, we exposed progenitors under dorsal and ventral spinal cord conditions to SB431542 from Day 5 onwards (Figure 6A). This treatment did not result in a change in the proportion of progenitors expressing Pax3 or Nkx2.2, suggesting it does not strongly affect the dorsal-ventral identity of neural progenitors (Figure 6C), but caused a significant delay in the induction of the late marker Nfia in neural progenitors and the expression of the intermediate and late-born markers Zfhx3, Nfia and Neurod2 in neurons under dorsal and ventral conditions (Figure 6D,E). To further investigate the consequences of TGFβ pathway inhibition on the temporal patterning of neural progenitors, we additionally assayed the consequences of ectopic TGFβ pathway activation and inhibition on the expression of the early genes *Lin28a, Lin28b* and the late genes *Sox9, Nfia, Nfib* and *Nfix* by RT-qPCR (Figure 6F). This analysis revealed a faster downregulation of early progenitor and earlier induction of late progenitor markers upon exposure to 2ng/ml TGFβ2 ligand (Figure 6G), while the opposite was true when the TGFβ pathway was inhibited using 10 µM SB431542 (Figure 6H). Together, these experiments demonstrate that TGFβ signalling controls the speed of progenitor maturation and the timing of temporal TF expression in in-vitro generated spinal cord neurons.

### Nfia and Nfib are required for the efficient generation of late-born spinal cord neurons

Nfi TFs are best known for promoting the switch from neurogenic to gliogenic progenitors (Deneen et al., 2006; Kang et al., 2012; Matuzelski et al., 2017; Tchieu et al., 2019). However, the expression of Nfia and Nfib in progenitors in the mouse spinal cord commences between e10.5 and e11.5, approximately 2 days before neurogenesis ceases and gliogenesis starts. Furthermore, these TFs are expressed in late-born neurons in the developing midbrain, hindbrain and spinal cord (Delile et al., 2019; Tiklová et al., 2019) (Figure 2D,I). In the retina, Nfi TFs are required for the specification of Müller glia and, importantly, bipolar cells, a late-born neuronal subtype (Clark et al., 2019). Nfi TFs also play important roles during the generation and postmitotic maturation of cerebellar granule neurons (Ding et al., 2013; Harris et al., 2015; Wang et al., 2010). These findings raise the possibility that Nfi TFs are also required for the specification of late-born neuronal subtypes in other parts of the central nervous system. To test this possibility, we generated an *Nfia*; *Nfib* double-mutant ES cell line by CRISPR/Cas9-induced non-homologous end joining. Because Nfia and Nfib act redundantly during the induction of gliogenesis in the spinal cord and the formation of bipolar cells and Müller glia in the retina (Clark et al., 2019; Deneen et al., 2006), we decided to directly focus on analysing the double mutant to rule out any potential redundancy between both genes. Electroporations of guide RNAs targeting the 2^nd^ coding exons of both genes resulted in double-heterozygous frameshift deletions of 20 and 11 base pairs in *Nfia* and 10 and 8 base pairs in *Nfib* (Figure S7A,B). Immunofluorescence assays of dorsal differentiations at Day 10 of differentiation, a timepoint when both proteins are normally detected at high-levels in progenitor nuclei in control differentiations, confirmed the absence of both proteins (Figure S7C,D).

Both in-vitro and in the developing spinal cord, Neurod2-positive neurons are born after Nfia and Nfib expression commenced in progenitors (Delile et al., 2019) (compare Figures 4B and 5E). Motif analysis (Fornes et al., 2020) revealed multiple Nfi motifs in close proximity to the *Neurod2* gene and analysis of recently published Nfia, Nfib and Nfix ChIP-seq datasets from the murine cerebellum (Fraser et al., 2020) confirmed binding of all 3 TFs to these sites (Figure 7A). We thus focused on assaying Neurod2 expression to determine the importance of Nfia and Nfib for the generation of late-born neurons in our in-vitro cultures. As our previous characterizations revealed the highest proportion of Neurod2-positive neurons are generated in ventral differentiations (Figure 4B), we focused on this condition. Characterizing the proportion of neurons expressing Neurod2 by flow cytometry revealed a marked reduction in the percentage of Neurod2-positive neurons in *Nfia; Nfib* double mutants (Figure 7B-D). Taken together, these data suggest that Nfia and Nfib are required for the expression of Neurod2 in late-born neurons in the spinal cord and support a model in which the specification of late-born neuronal subtypes is tightly coupled to the signals and transcriptional programs that mediate the switch from neuro-to gliogenesis throughout the nervous system.

**Figure 7:**
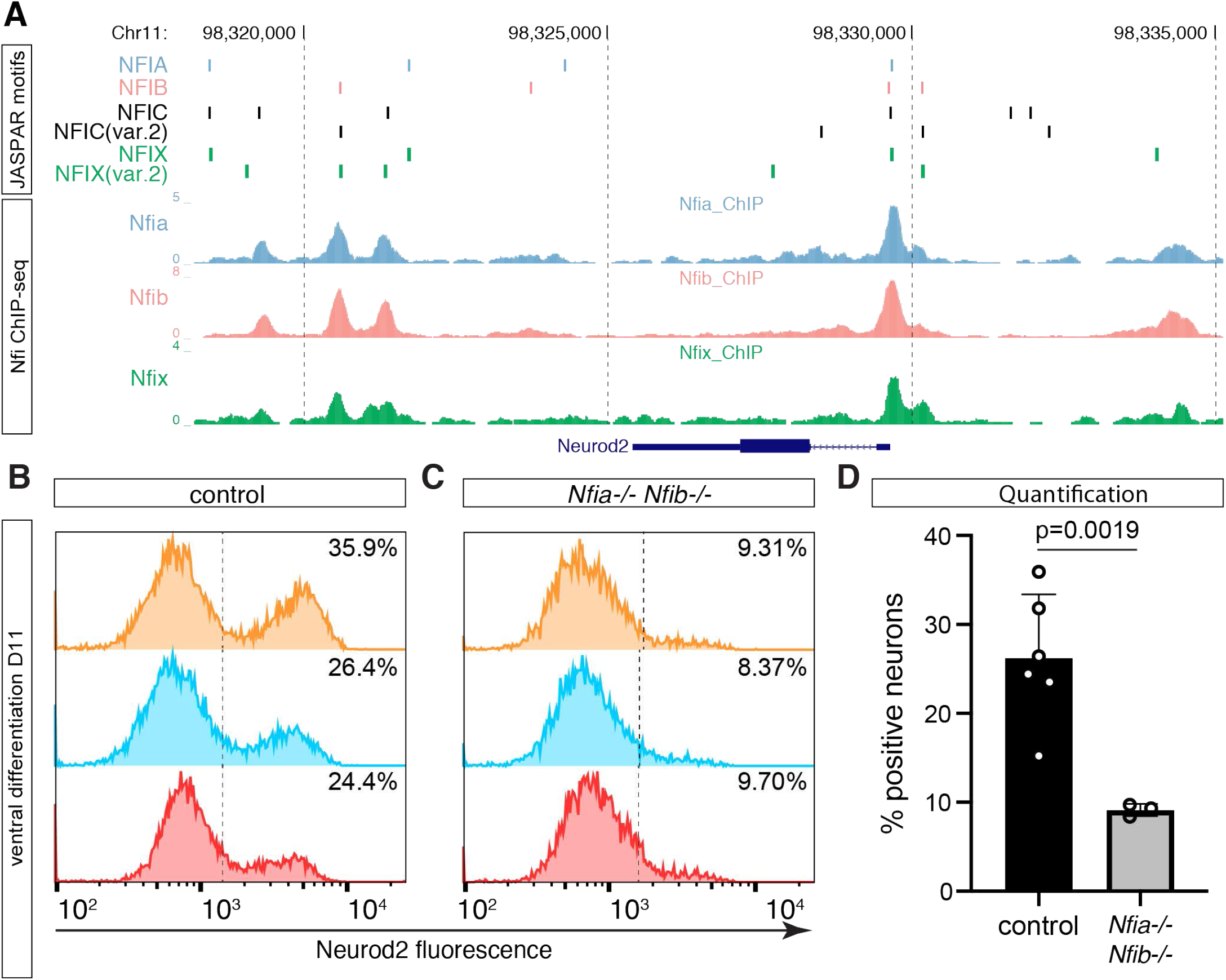
Nfia and Nfib are required for the efficient generation of late-born Neurod2 neurons (see also Figure S6) (A) Analysis of the Neurod2 locus using JASPAR (Fornes et al., 2020) identifies multiple Nfi transcription factor binding motifs (top). Analysis of Nfia, Nfib and Nfix ChIP-Seq data from the mouse cerebellum (Fraser et al., 2020) confirms binding of all three TFs to these motifs (bottom). (B,C) Neurod2 intensity histograms in control (B) and *Nfia*; *Nfib* double mutant (C) neurons at D11 in ventral conditions. Dashed lines indicate the applied thresholds above which neurons were counted as Neurod2 positive. (D) Quantification of the percentage of Neurod2-positive neurons at D11 in control and *Nfia; Nfib* double mutants differentiated in ventral conditions reveals a strong reduction of Neurod2 neurons in the absence of Nfia and Nfib. (n=6 for control and n=3 for Nfia; Nfib double mutants). Significance was assessed by unpaired t-test with Welch’s correction.

## Discussion

### Neuronal diversity from the superposition of spatial and temporal patterning programs

The recent advent of single cell sequencing technologies, such as RNA sequencing, has enabled the profiling of cell-type diversity at unprecedented scale (Briscoe and Marín, 2020). Especially in the nervous system, this has led to the discovery of a much greater complexity and molecular heterogeneity of cell types than previously anticipated, raising the question of how this diversity arises. The superposition of multiple patterning systems that act along different spatiotemporal axes provides a solution to this problem as it enables the combinatorial specification and organisation of cell types using relatively simple patterning schemes (Erclik et al., 2017). Here we provide evidence of a temporal patterning programme, operating in parallel to spatial mechanisms, throughout the vertebrate nervous system.

The same temporal sequence of TF expression is observed in forebrain, midbrain, hindbrain, spinal cord, retina, and in ES cell-derived neurons with various axial identities. We also defined a temporal patterning program within neural progenitors and demonstrate that TGFβ signalling controls the pace of the program. Moreover, we find that the TFs Nfia and Nfib, typically considered markers of gliogenic potential, are required for the generation of late-born neurons in an in-vitro model of spinal cord development. Taken together, our results suggest that the conserved temporal patterning of progenitors and neurons mediated by TGFβ signalling contributes to the generation of neuronal diversity in large parts of the nervous system, including disease-relevant cell types such as midbrain dopaminergic neurons.

The temporal program functions in parallel to the well-established spatial patterning of the dorsal-ventral and anterior-posterior axes of the neural tube, thus enabling the generation of a combinatorially increasing number of neuronal subsets from the superposition of a limited number of TFs that delineate specific spatial and temporal identities. This mechanism could be extended further. For example, the temporal patterning of Drosophila medulla neuroblasts is defined by the sequential expression of 5 TFs, however, the expression of these temporal TFs in aging neuroblasts is not mutually exclusive – instead there are periods of co-expression of sequentially expressed TFs (Li et al., 2013). Similar observations have been made in the neuroblast lineages in the Drosophila embryo and mushroom body (Averbukh et al., 2018; Liu et al., 2019). Such co-expression of temporal TFs has been proposed to designate additional temporal windows during which further neuronal subtypes are generated (Averbukh et al., 2018; Li et al., 2013). Our data, so far, do not support such a model in the spinal cord, as the expression of the different pairs of temporal TFs we analysed in spinal cord neurons were mutually exclusive (Figure S2). We note, however, that the respective temporal identities are defined by co-expression of multiple orthologous TFs and we analysed a limited number of TF pairs. Moreover, characterization of spinal V1 interneuron diversity has revealed differential expression of Onecut1 and Onecut2 in some V1 subtypes, some of which also expressed Zfhx4 (Gabitto et al., 2016). Taken together, these observations raise the possibility that the temporal TF code could further diversify the number of neurons generated in each domain based on the combinatorial co-expression of distinct pairs of temporal TFs. Future experiments are required to test this hypothesis.

Concomitantly with neurons, neural progenitors throughout the vertebrate nervous system undergo a temporal patterning program (Figure 5A,B). Components of this program, including Sox9 and Nfia/b, have previously been implicated in the transition from neurogenesis to gliogenesis (Deneen et al., 2006; Kang et al., 2012; Namihira et al., 2009; Scott et al., 2010; Stolt et al., 2003). However, the expression of these factors precedes the onset of gliogenesis. The expression of Sox9 in neural progenitors coincides with the switch from early Onecut2-positive to intermediate Pou2f2 and Zfhx3-positive neurons and the induction of Nfia/b correlates with the later transition. Moreover, the loss of generation of late neuronal subtypes in neural progenitors lacking Nfia/b is consistent with the involvement of these TFs in the neuronal temporal program as well as the gliogenic switch (Figure 7). This raises the possibility that the transition of neural progenitors from exclusively neurogenesis to subsequent gliogenesis is part of the same temporal patterning program operating in the nervous system. This would be analogous to the temporal program in Drosophila neuroblasts which also controls the identity of neurons and glia cells.

### TGFβ signalling regulates temporal patterning in the nervous system

Our results indicate that the TGFβ pathway is an important regulator of the pace of progenitor maturation and the timing of temporal TF expression in neurons. This is in agreement with previous findings. In the hindbrain, TGFβ2 signalling controls the timing of the switch from MNs to serotonergic neurons by repressing the TF Phox2b in neural progenitors (Dias et al., 2014). In addition, another TGFβ family member, Gdf11, controls the timing of retinal ganglion cell specification in the vertebrate retina, the timing of MN subtype specification, and the switch from dI5 to late-born dIL_B_ neurons in the spinal cord (Kim et al., 2005; Shi and Liu, 2011). The connection between these roles of TGFβ signalling and its role in directing the temporal patterning programs of progenitors and neurons is currently unclear but, taken together, the results implicate multiple ligands of the TGFβ family in controlling the temporal patterning of the mammalian nervous system. Notably, Activin signalling is involved in controlling the timing of fate switches in the Drosophila mushroom body and, similar to observations in vertebrates, inhibition of Activin signalling results in a delay of temporal fate progression in this system (Rossi and Desplan, 2020). These findings suggest a deep evolutionary origin for the role of the TGFβ pathway in controlling temporal patterning and the diversification of cell types in the developing nervous system bilaterians.

The timing of switches in temporal TF expression occur at approximately similar times throughout the developing nervous system and during the in-vitro differentiation of neurons with different axial and dorsal-ventral identities. This raises the question how signals, from locally secreted sources, achieve an apparently globally synchronised effect and what the source of TGFβ might be in the in-vitro differentiations. A solution to this puzzle could be that Gdf11 and related ligands are expressed in new-born neurons in the spinal cord (Shi and Liu, 2011). Notably, analysis of Gdf11 expression suggests that this expression pattern is preserved in the hindbrain and midbrain. Such a model, in which the temporal progression of progenitors is coupled to a ligand secreted by neurons that signals back to progenitors has the advantage that it provides a means to ensure that the correct proportion of neurons with a specific temporal identity are produced before progenitors switch to the next phase. A prediction of such a model is that local increases in neurogenesis would lead to a local acceleration of temporal patterning in progenitors. Indeed, several genes involved in the onset of gliogenesis, such as *Sox9, Nfia* and *Fgfr3*, are first expressed in the ventral spinal cord (Deneen et al., 2006; Kang et al., 2012; Stolt et al., 2003), where MNs differentiate at higher rate at early developmental stages (Kicheva et al., 2014; Novitch et al., 2001). Further experiments that explore the connection between Gdf11, neurogenesis rate and temporal patterning are required to test this hypothesis.

The data show that the temporal pattern of both neurons and progenitors continues to advance in the absence of TGFβ pathway activity. This is consistent with observations from the ventral hindbrain, where ablation of Tgfbr1 delays but does not abrogate the switch from MNs to serotonergic neurons and in Gdf11 mutants in the spinal cord, where the onset of oligodendrocyte formation is delayed but not prevented (Dias et al., 2014, 2020; Shi and Liu, 2011). Together this suggests that other extrinsic signals, or cell-intrinsic timers, must exist that promote temporal progression. A potential candidate signal that may oppose the activity of TGFβ is retinoic acid (RA), which has been shown to drive the generation of Onecut-positive Renshaw cells in an in-vitro model of V1 subtype diversity (Hoang et al., 2018). Furthermore, the rate-limiting enzyme for RA synthesis is down-regulated in somites, adjacent to the neural tube, between e9.5 and e10.5 (Niederreither et al., 1997), coinciding with the switch from Onecut to Zfhx3 positive neurons. In addition, several pathways, such as Neuregulins, Notch, FGF and JAK/STAT have been shown to promote gliogenesis (Miller and Gauthier, 2007; Namihira et al., 2009; Vartanian et al., 1999). Given the pivotal role of Nfi TFs in this process, one or more of these signals may promote the acquisition of a late Nfi-positive progenitor identity. The genetic and experimental accessibility of in-vitro models will allow these possibilities to be tested.

### Temporal patterning of in-vitro generated neurons

In-vitro generated neurons are widely used for disease modelling and have the potential to offer novel therapeutic avenues to tackle nervous system injuries and neurodegenerative diseases (Fischer et al., 2020; Sances et al., 2016; Tao and Zhang, 2016). A better understanding of the molecular mechanisms responsible for neuronal diversity contributes to the rational design of in-vitro differentiation protocols to generate cell types best-suited for such applications. Our work demonstrates that the temporal patterning of neurons and progenitors is conserved in-vitro, providing a new dimension for assessing the identity of progenitors and neurons obtained in culture. Furthermore, the observation that manipulating TGFβ signalling can accelerate or slow-down the progression of temporal patterning opens up the possibility to use such perturbations to increase the yield of progenitors and neurons with desired spatial and temporal identities.

Many applications of in-vitro generated neurons and progenitors require large numbers of cells with defined identities. These are often generated by expanding progenitors using treatments with signals such as EGF and/or FGF before exposing the resulting progenitor populations to differentiation stimuli. Such prolonged expansion phases might result in the preferential generation of neurons with late temporal identities. This might be at least partially counter-acted by the incorporation of TGFβ-pathway inhibitors. Indeed, treatment with SB431542 in combination with other small molecules has been demonstrated to enable long-term self-renewal of neural stem cells for more than 30 passages (Li et al., 2011). Another promising approach to generate neurons with defined identities is reprogramming of pluripotent cells or somatic cells, such as fibroblasts or astrocytes, using specific cell-fate converting cocktails of transcription regulators. Notably, the reprogramming of ES cells to different types of neurons resulted in expression of Onecut TFs (Aydin et al., 2019; Mazzoni et al., 2013), suggesting such approaches might preferentially generate the earliest temporal identities. The addition of temporal TFs that define later stages of the differentiation program to these reprogramming cocktails might expand the toolbox for the efficient generation of a wider-range of neuronal subtypes with desired temporal identities for in-vitro disease modelling and future clinical applications.

## Experimental Procedures

### Animal Welfare

Animal experiments were performed under UK Home Office project licenses (PD415DD17) within the conditions of the Animal (Scientific Procedures) Act 1986. All experiments were conducted using outbred UKCrl:CD1 (ICR) (Charles River) mice.

### Immunofluorescent staining and microscopy

Embryos were fixed at the indicated stages in 4% PFA (Thermo Fisher Scientific) in PBS on ice, cryoprotected and dissected in 15% ice-cold sucrose in 0.12M PB buffer, embedded in gelatine and 14 µm sections taken. In-vitro generated cells were fixed for 15 minutes in 4% PFA in PBS at 4 degrees. 30 minutes blocking and primary antibody incubation over night at 4 degrees was performed using PBS + 0.1% Triton (PBS-T) + 1% BSA. A complete list of antibodies is available in Table S1. The next day, samples were washed 3x 30 minutes in PBS-T and incubated with secondary antibodies in PBS-T + 1% BSA for 1 hour at room temperature. Secondary antibodies used throughout the study were raised in donkey (Life Technologies, Jackson Immunoresearch). Alexa488 and Alexa568-conjugated secondary antibodies were used at 1:1000, Alexa647-conjugated antibodies at 1:500. Samples were washed 3 more times in PBS-T and then mounted in Prolong Antifade (Molecular Probes).

For EdU-labelling, mice were intraperitonially injected with 3 µl/gramm body weight EdU diluted in PBS at the indicated stages. EdU was detected using Alexa647 Click-iT EdU Imaging Kit (Invitrogen C10340) according to the manufacturer’s specifications. At least 4 sections from different animals were analysed for each timepoint.

Stainings of in-vitro differentiations were acquired on a Zeiss Imager.Z2 microscope equipped with an Apotome.2 structured illumination module and a 20× air objective (NA=0.75). Cryo-sections were imaged using a Leica SP8 equipped with a 40x oil PL APO CS2 objective (NA=1.30). Tissue sections were tiled using 10% overlap between adjacent tiles and merged using LAS X software.

### Image analysis

Image analysis was performed in Fiji (http://fiji.sc/Fiji) and Python3.7 (http://www.python.org). e13.5 mouse neural tube transverse sections were manually cropped using Fiji and then processed using a custom Python pipeline. Cell nuclei were segmented using an adaptive threshold and watershed algorithm on the DAPI channel. Parameters for proper segmentation and filtering were manually optimized for each set of images. Segmented objects were further filtered based on area to fit the expected nuclei dimensions. Neuronal nuclei were distinguished from those of progenitors either by presence of the neuronal marker HuC or absence of Sox2 staining. For each neuronal nucleus the mean intensity of the temporal TFs and EdU was then calculated.

Data analysis and plotting was performed in R (https://www.r-project.org). For each section, intensities in nuclei were first normalized between 0 and 1. To remove outliers 0.3% of the brightest and dimmest objects were discarded. Objects were counted as positive for EdU or expression of temporal TFs if their normalized intensity was greater than 0.25. Percentage of EdU-positive nuclei expressing temporal TFs was then plotted using ggplot2 (Wickham, 2016).

### ESC culture and differentiation

HM1 mouse ESCs (Thermo Fisher Scientific) were maintained and differentiated as described previously (Gouti et al., 2014; Metzis et al., 2018; Sagner et al., 2018). In brief, ESCs were maintained on a layer of mitotically inactivated mouse embryonic fibroblast (feeders) in ES cell medium + 1,000 U/ml LIF. For differentiation, ESCs were dissociated using 0.05% Trypsin (Gibco). Feeder cells were removed by replating cells for 25 minutes on a tissue culture plate. 60-80,000 cells were plated onto 0.1% Gelatin (Sigma) coated 35 mm CellBIND dishes (Corning) into N2B27 medium + 10 ng/ml bFGF. Differentiation protocols for progenitors and neurons with different axial and dorsal-ventral identities are shown in Figure 4A. Differentiation protocols for activation and inhibition of the TGFß pathway using TGFß2 (R&D Systems) or 10 µM SB431542 (Tocris) respectively are shown in Figure 6A,F. For midbrain differentiation cells were kept in N2B27 medium with addition of 10 ng/ml bFGF until Day 3. To generate hindbrain identity 100 nM RA (Sigma) and 500 nM SAG (Calbiochem) were supplemented together at Day 3 and 4. For spinal cord differentiations, cells were exposed to 5 μM CHIR99021 (Axon) between days 2 and 3 and then supplemented with 100 nM RA (Sigma) until day 5. For ventral differentiations, cells were additionally exposed to 500 nM SAG (Calbiochem) from days 3 to 9.

### Generation of *Nfia; Nfib* double mutant ESCs

For generation of *Nfia; Nfib* double mutant ESCs CRISPR guide RNAs were cloned into pX459 plasmid obtained from Addgene (# 62988), according to Ran et al., 2013. ESCs were electroporated using Nucleofector II (Amaxa) and mouse ESC Nucleofector kit (Lonza). Afterwards, cells were replated onto 10-cm CellBind plates (Corning) and maintained in 2i medium + LIF. For selection, cells were first treated with 1.5 μg/ml Puromycin (Sigma) for two days and afterwards maintained in 2i medium + LIF until colonies were clearly visible. Individual colonies were picked using a 2-μl pipette, dissociated in 0.25% Trypsin (Gibco), and replated onto feeder cells in ES-medium + 1,000 U/ml LIF in a 96-well plate. Mutations in *Nfia* and *Nfib* were analyzed by PCR over the targeted regions and verified by Sanger sequencing. Overlapping peaks arising from heterozygous indels were deconvolved using CRISP-ID (Dehairs et al., 2016) (Figure S6A,B). Loss of Nfia and Nfib protein was further confirmed by immunofluorescent staining at day 10 of the differentiation (Figure S6C,D).

### Flow Cytometry

In-vitro differentiations were dissociated at the indicated timepoints using 0.05% Trypsin (Gibco). Live/Dead cell staining was performed using LIVE/DEAD Fixable Near-IR Dead Cell Stain Kit (Invitrogen) for 30 minutes on ice. Immediately afterwards cells were spun-down for 2 min at 1000xg and fixed in 4% PFA for 12 minutes on ice. Fixed cells were spun-down, resuspended in 500 µl PBS, and stored at 4 degrees for up to 2 weeks.

For staining 1.5 – 2 million cells were used. Cells were spun-down and incubated with antibodies in PBS-T + 1% BSA. If primary and secondary antibodies were used, cells were incubated in primary antibody solution over night. Directly-conjugated antibodies or secondary antibodies were applied for 1 hour at room temperature. A complete list of antibodies used for flow cytometry is supplied in Table S2. Flow cytometry analysis was performed using BD LSR Fortessa analyzers (BD Biosciences). Data analysis was performed using FlowJo (v10.4.1) and plotted using Graphpad Prism 7. The general gating strategy is outlined in Figure S4B. Progenitor and neuronal cell populations were discriminated based on Sox2 and Tubb3 antibody staining (Figure S4B). Percentages of Onecut2, Neurod2 and Zfhx3-positive neurons were calculated by applying a threshold at which 1-2% of Sox2+ progenitors in the same sample were counted as positive. Percentage of Nfia-positive neurons and progenitors was determined using a global threshold for all datasets. Data was plotted and statistical analysis performed in GraphPad Prism 8. Graphs throughout the manuscript show means ± standard deviation of all conducted replicates. Statistical significance was assessed using unpaired t-tests. A summary of the percentage of positive cells, replicate number and p-values is provided in Table S3. Significance values throughout the manuscript are indicated by p<0.001 = ***, p<0.01 = **, p<0.05 = *.

### RT-qPCR

Total RNA was isolated from cells at the indicated time points using Qiagen RNeasy kit according to the manufacturers instructions. Genomic DNA was removed by digestion with DNase I (Qiagen). cDNA synthesis was performed using SuperScript III (Invitrogen) and random hexamers. qPCR was performed using PowerUp SYBR Green Master Mix (Thermo Fisher Scientific) using 7900HT Fast Real time PCR (Applied Biosystems), QuantStudio 5 or QuantStudio 12K Flex Real-Time PCR Systems (Thermo Fisher Scientific). qPCR primers were designed using NCBI tool Primer BLAST and are listed in Table S4. All experiments were conducted at least in biological triplicates for each timepoint analysed. Expression values were normalized to ß-actin. Data was plotted and statistical analysis performed in GraphPad Prism 8. Graphs throughout the manuscript show means ± standard-deviation of all replicates.

### scRNAseq data analysis

scRNAseq analysis was performed using R-Studio v1.2.1335 using R v3.5.2. A complete R script decribing the scRNAseq analysis performed in this paper is available at https://github.com/sagnera/tTF_paper_2020

### Differential gene expression analysis in spinal cord neural progenitors

scRNAseq data from e9.5-e13.5 spinal cord neural progenitors including subtype annotations were obtained from Delile et al., 2019. dp6 progenitors were excluded from this analysis due to low numbers in the dataset. For each progenitor domain, differential gene expression between progenitors from different embyronic days was performed using Seurat v3.1.4 (Stuart et al., 2019) using the “FindAllMarkers” function with settings min.pct = 0.25 and logfc.threshold = 0.25. Only genes detected in more than 7 progenitor domains were retained. TFs were identified based on a list of TFs encoded in the mouse genome obtained from AnimalTFDB3.0 (Hu et al., 2019). Heatmaps in Figure 5A show log-scaled and z-scored gene expression.

### Analysis of temporal TFs in the mouse retina

scRNAseq of the developing retina (Clark et al., 2019) was downloaded from https://github.com/gofflab/developing_mouse_retina_scRNASeq and imported into Seurat v3.1.4 (Stuart et al., 2019). Cells were filtered based on age (e14, e16, e18 and P0), cell type (RPCs, Neurogenic Cells, Photoreceptor Precursors, Cones, Rods, Retinal Ganglion Cells, Amacrine Cells, Horizontal Cells), number of reads in each cell (nFeature > 800 and nFeature < 6000) and percentage of reads in mitochondrial genes (percent.mt < 6). Only cells annotated as Horizontal Cells, Amacrine Cells, Retinal Ganglion Cells, Rods and Cones were used for the time-stratified heatmap of temporal TF expression.

### Expression dynamics of temporal TFs in the fore-, mid- and hindbrain

Annotated scRNAseq data from the developing fore-, mid- and hindbrain was downloaded from mousebrain.org (Manno et al., 2020). Cells were assigned fore-, mid- and hindbrain identity based on the “Tissue” column of the provided loom file. To account for the different sequencing depths between cells, readcounts were normalized by multiplying the counts in each cell with 10,000 divided by the total number of UMIs in this cell. Mean expression and ratio of expressing cells for the indicated temporal TFs and regions were calculated in R. Data was plotted using ggplot2 (Wickham, 2016). Heatmaps in Figure 5B show log-scaled and z-scored gene expression.

### Comparison with in-vitro RNAseq data

RNAseq data from D3-D10 ventral spinal cord differentiations (Rayon et al., 2020) (GSE140748) was used. Gene expression per timepoint was averaged over all 3 provided replicates. Only data from full days of differentiation (D3, D4, D5, D6, D7, D8, D9, D10; D0-D7 in the provided data files) was used for further analysis. Heatmaps in Figure 5C show log-scaled and z-scored gene expression.

### Alignment of Nfia/b/x ChIP-seq data

Nfia/b/x ChIP-seq data from Fraser et al., 2020 was downloaded from the GEO database (GSE146793) and aligned to mm10 using the nf-core ChIP-seq pipeline v1.1.0 (Ewels et al., 2020).

## Acknowledgements

We thank all members of the Briscoe lab for help, advice, reagents and critical feedback. We acknowledge scientific support by the Crick Science and Technology platforms, in particular the Biological Research Facility, Equipment Park, Flow Cytometry and Light Microscopy facilities. We thank M.J. Delás for help with flow cytometry; Thomas Müller, Carmen Birchmeier and Siew-Lan Ang for kindly sharing antibodies; and Nancy Papalopulu, Tiago Rito and François Guillemot for comments on the manuscript.

## Author Contributions

Conceptualization: A.S., J.B.; Methodology: A.S., I.Z., T.W., M.M., J.B.; Software: A.S., J.L.; Investigation: A.S., I.Z., T.W., M.M.; Resources: T.W.; Writing - original draft: A.S., J.B.; Writing - review & editing: A.S., I.Z., T.W., J.L., J.B.; Supervision: A.S., J.B.; Project administration: J.B.; Funding acquisition: A.S., J.B.

## Supplemental Figure legends

**Figure S1.**
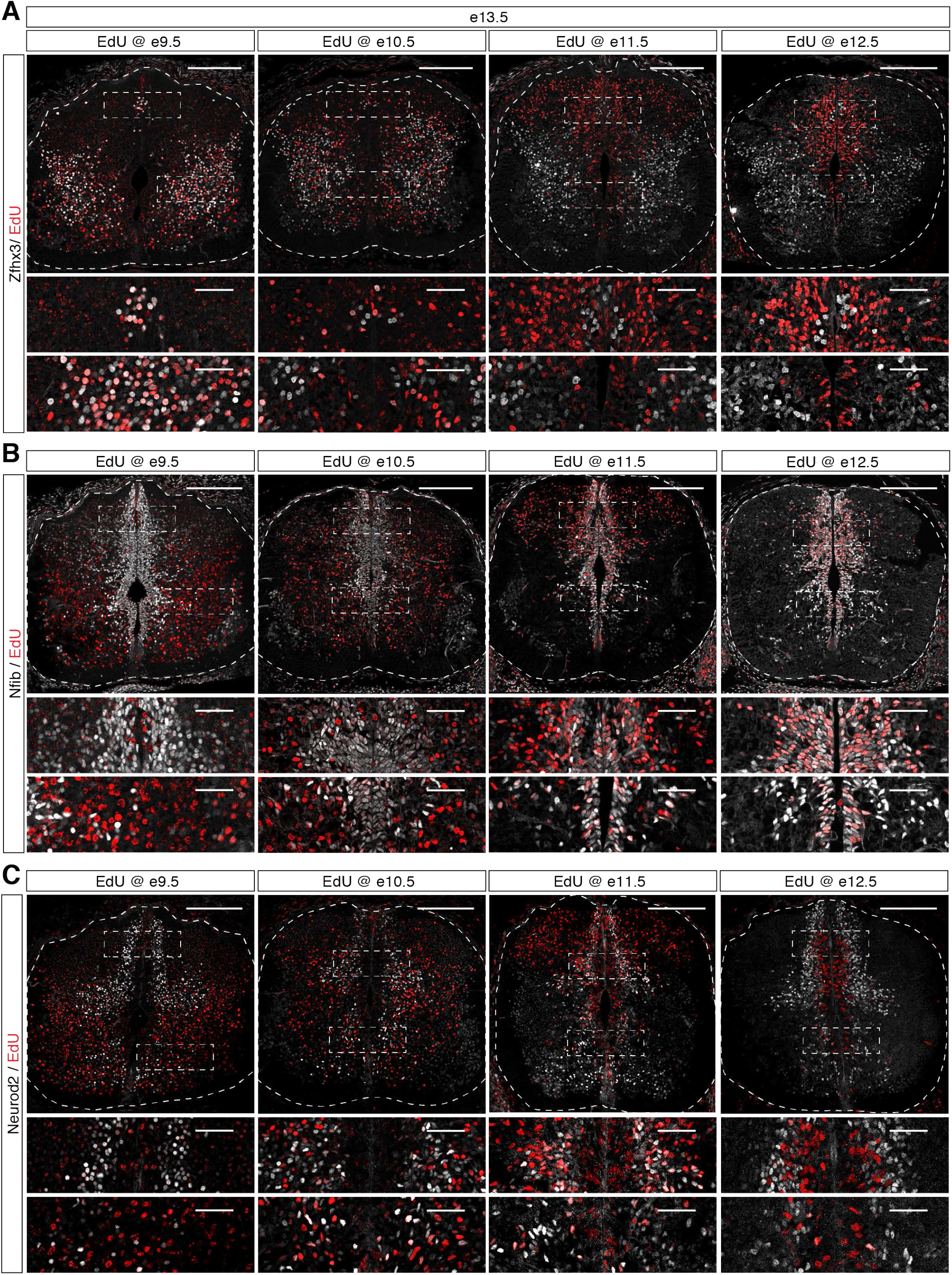
Related to Figure 1: Complete time course of colocalization between temporal TFs and EdU administered at different timepoints. (A-C) Colocalization between Zfhx3 (A), Nfib (B), Neurod2 (C) and EdU administered at e9.5, e10.5, e11.5 or e12.5 (from left to right) in e13.5 spinal cord sections. Scale bars in overview pictures = 200 µm, insets = 50 µm

**Figure S2.**
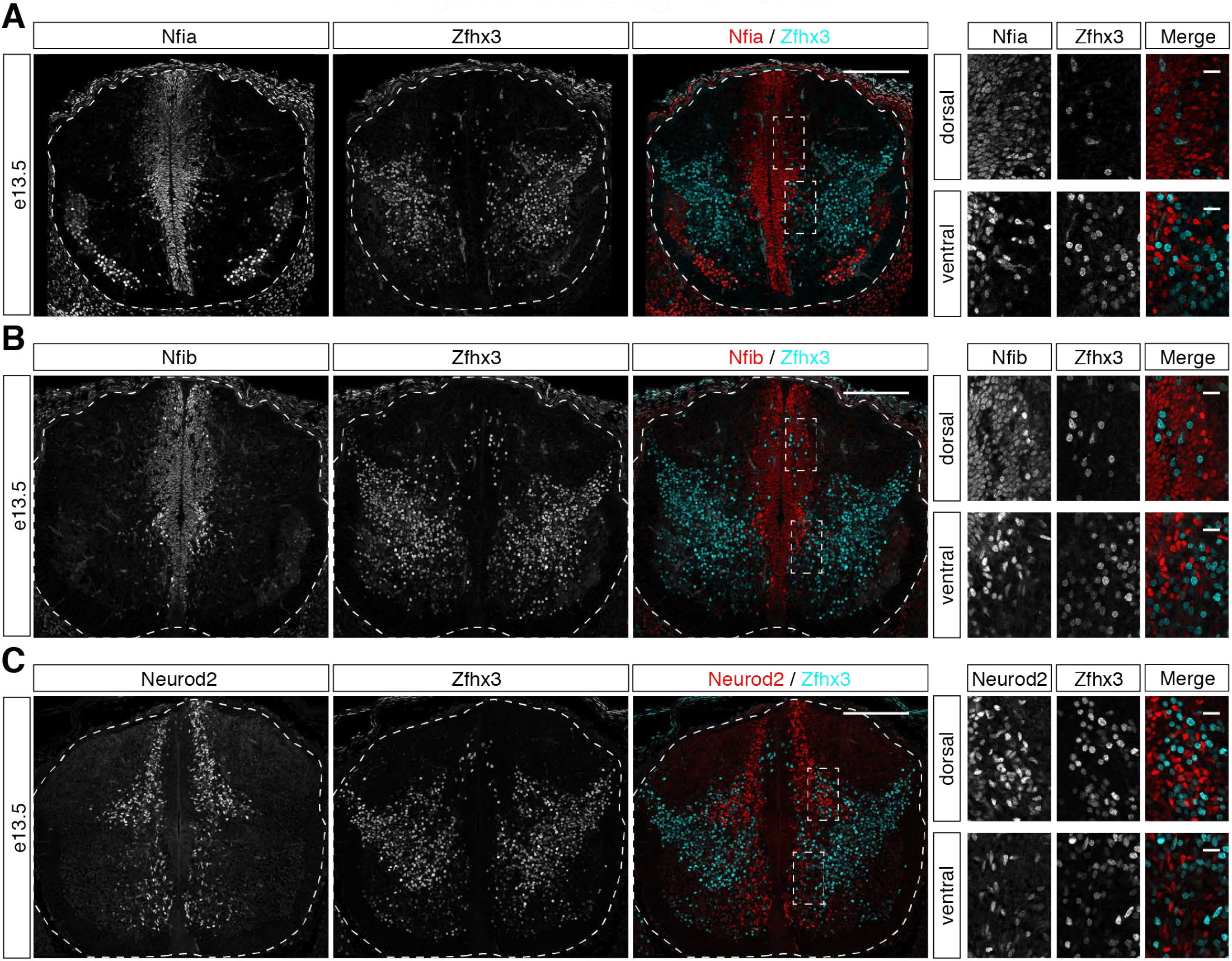
Related to Figure 1: Non-overlapping expression of temporal TFs in spinal cord neurons at e13.5. (A-B) Zfhx3 and Nfib (A) or Zfhx3 and Neurod2 (B) are expressed in mutually exclusive populations of neurons in the spinal cord. Scale bars in overview pictures = 200 µm, insets = 20 µm

**Figure S3.**
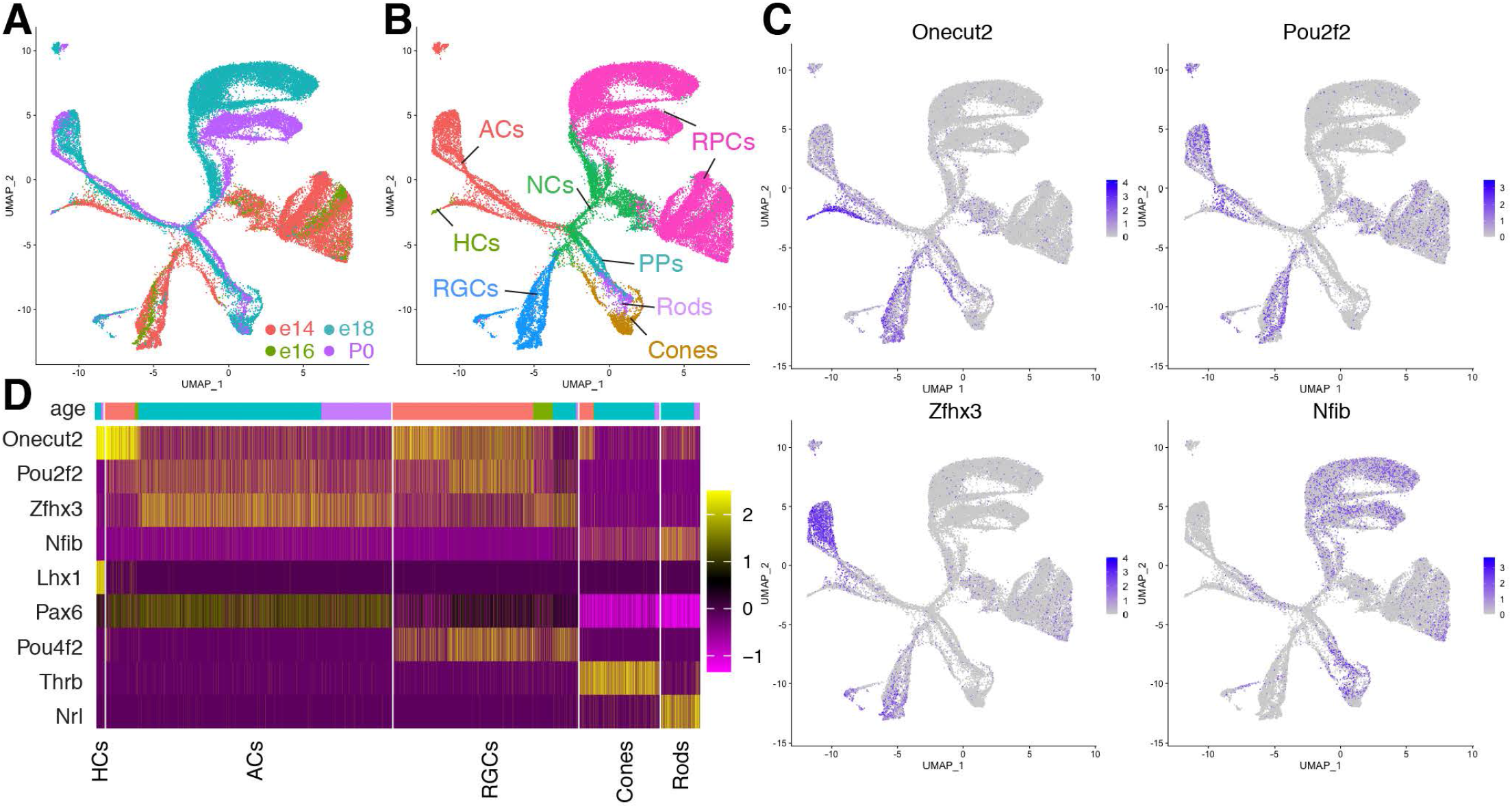
Related to Figure 2: Characterization of temporal TF expression in the developing retina. (A) UMAP representation of scRNAseq data from the developing mouse retina (Clark et al., 2019) color-coded by developmental stage. (B) Same UMAP-representation as (A) color-coded for cell identity (AC amacrine cells, HC horizontal cells, RGCs retinal ganglion cells, RPCs retinal progenitor cells, NCs neurogenic cells, PPs photoreceptor precursors) (C) Expression levels of Onecut2, Pou2f2, Zfhx3, and Nfib in individual cells (D) Heatmap indicating expression levels of the temporal TFs (Onecut2, Pou2f2, Zfhx3, and Nfib) and known marker genes (Lhx1, Pax6, Pou4f2, Thrb and Nrl) in different types of retinal neurons stratified by developmental age.

**Figure S4.**
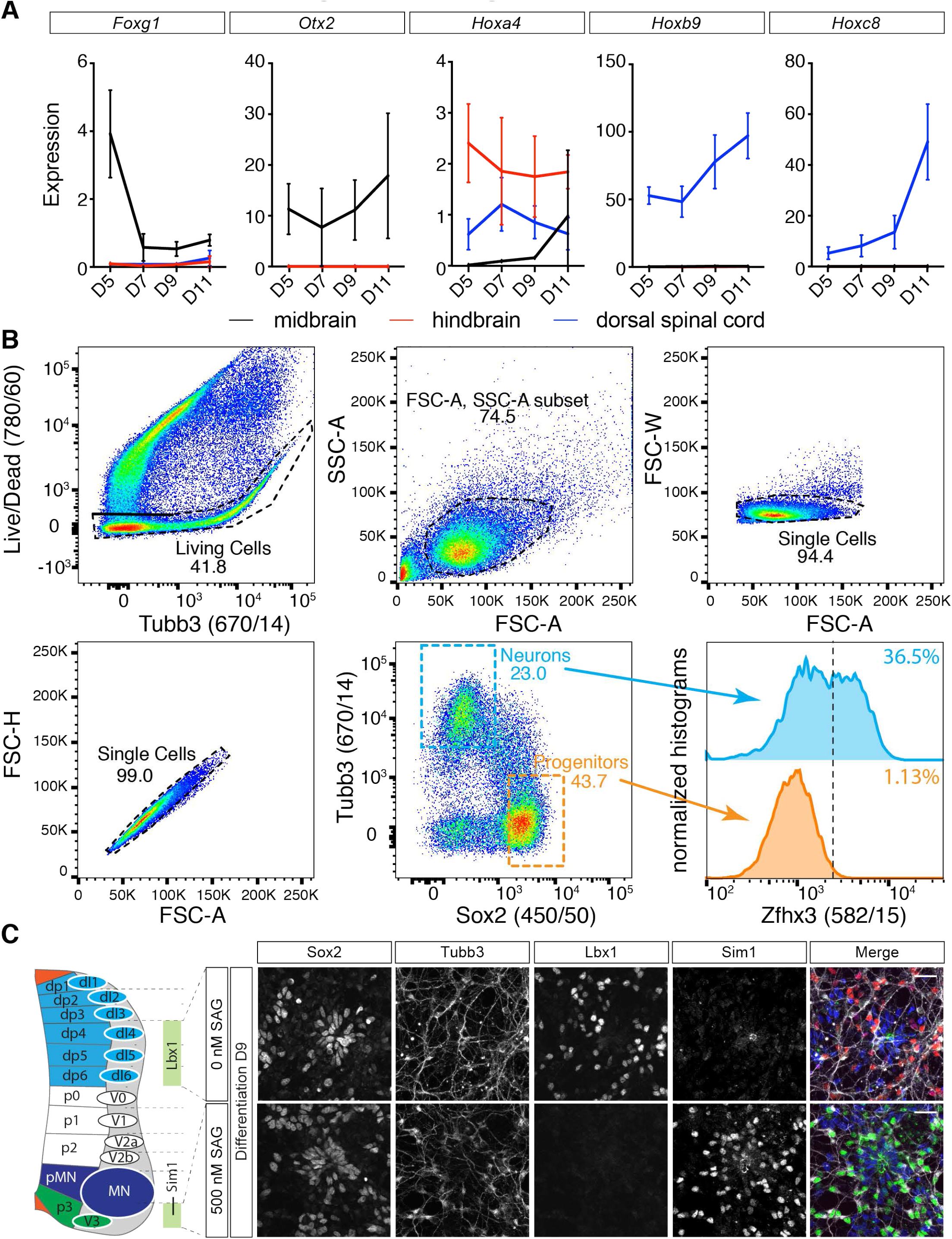
Related to Figure 4: Further characterization of the in-vitro differentiations. (A) RT-qPCR analysis of *Foxg1, Otx2, Hoxa4, Hoxb9* and *Hoxc8* reveals the generation of neurons and progenitors with different axial identities in the in-vitro differentiations (B) Gating strategy for the quantification of the expression of different markers in neurons and progenitors by flow cytometry. Living cells were identified based on Infrared Life/Dead stain. Gating on single cells was achieved using forward and side-scatter as indicated. Progenitors and neurons were discriminated based on the progenitor marker Sox2 and neuronal beta-tubulin (Tubb3). To quantify the proportion of neurons expressing Onecut2, Zfhx3 and Neurod2 an intensity threshold was applied to each sample that was exceeded by 1-2% of progenitors. The same threshold was then applied to neurons in the same sample and the percentage of neurons exceeding this threshold was counted as positive. As Nfia is expressed in neurons and progenitors, a global threshold was applied to quantify the proportion of neurons and progenitors expressing Nfia. (C) Characterization of dorsal and ventral spinal cord differentiations by immunostaining. Under dorsal conditions most neurons express the TF Lbx1, which is expressed in dI4-dI6 neurons generated in the intermediate dorsal part of the spinal cord. Under ventral conditions neurons express the V3 interneuron marker Sim1. Scale bars in C = 25 µm

**Figure S5.**
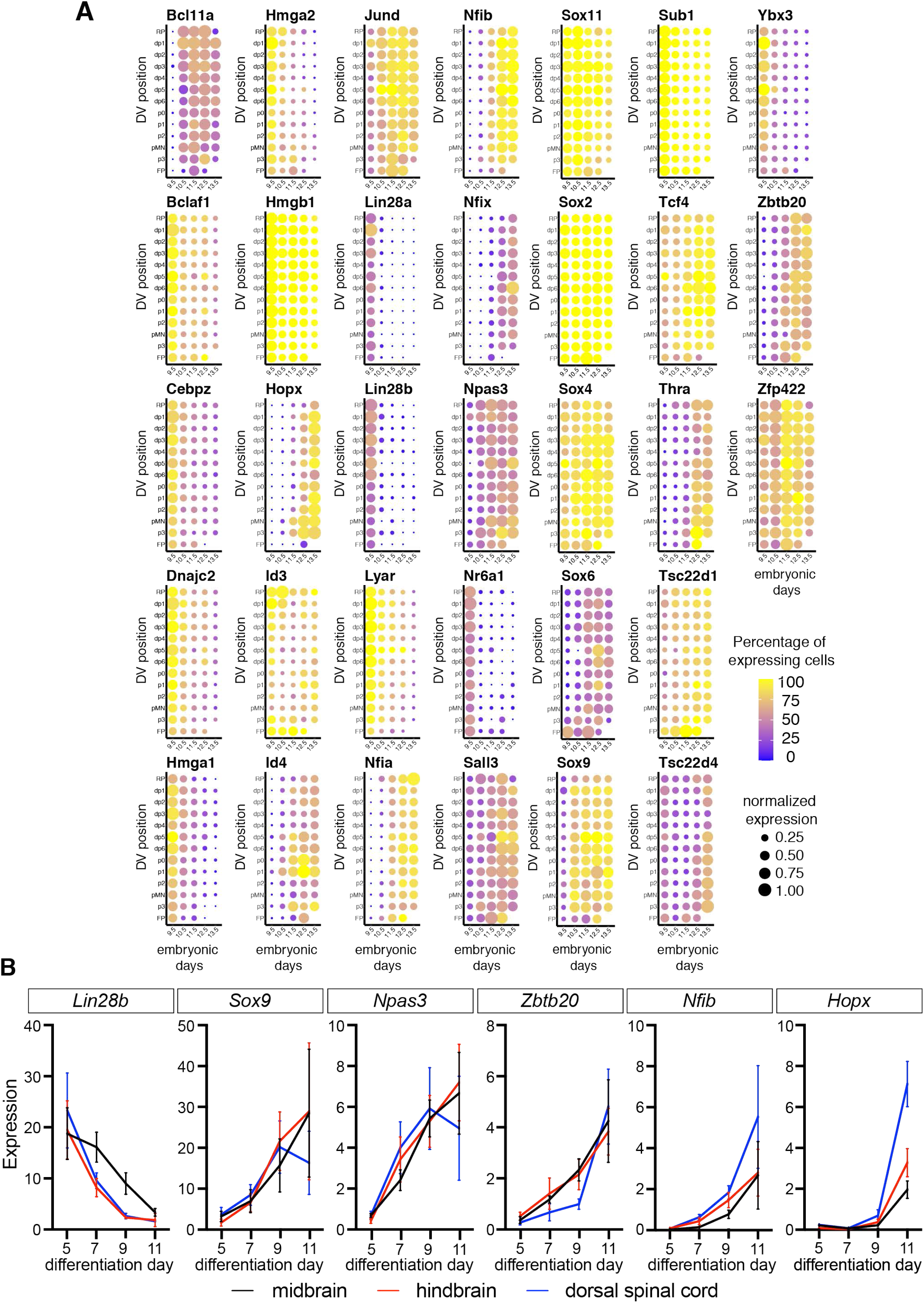
Related to Figure 5: Temporal patterning of neurons and progenitors. (A) Spatial and temporal expression of the 33 differentially expressed TFs during the neurogenic period in spinal cord neural progenitors (B) RT-qPCR analysis for *Lin28b, Sox9, Npas3, Zbtb20, Nfib* and *Hopx* from days 5-11 in in-vitro generated differentiations with different axial identities reveals that temporal patterning is conserved in-vitro.

**Figure S6.**
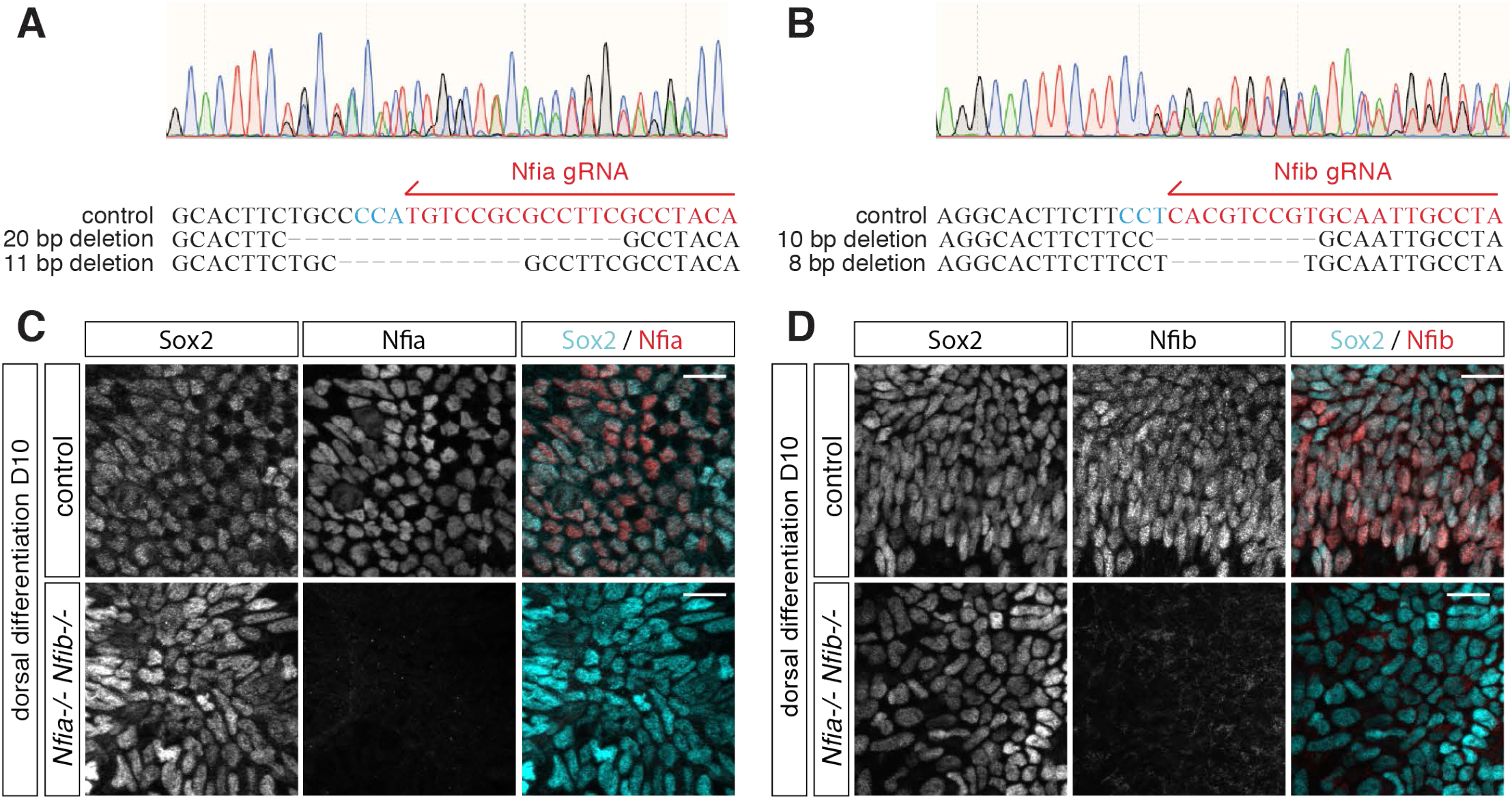
Related to Figure 7: Characterization of the *Nfia*; *Nfib* double mutant ES cell line. (A,B) Engineering of a *Nfia; Nfib* double mutant ES cell line by CRISPR/Cas9-mediated mutagenesis. Introduction of double heterozygous frameshift mutations in both genes was validated by Sanger sequencing. (C,D) Loss of Nfia (C) and Nfib (D) immunostaining in neural progenitors generated from *Nfia; Nfib* double mutant ES cells in dorsal differentiations at D10 Scale bars in C,D = 20 µm

## Supplemental Figure legends

**Table S1.** Related to Experimental Procedures: List of antibodies used for immunofluorescence.

| Antibody | Species | Company | Catalogue number | Dilution |
| --- | --- | --- | --- | --- |
| HuC | mouse | Molecular Probes | clone 16A11 | 1:250 |
| Lbx1 | guinea-pig | from Thomas Müller and Carmen Birchmeier | Müller et al. 2002 | 1:10000 |
| Lmx1b | guinea-pig | from Thomas Müller and Carmen Birchmeier | Müller et al. 2002 | 1:10000 |
| Neurod2 | rabbit | Abcam | ab104430 | 1:1000 |
| Nfia | rabbit | Atlas Antibodies | HPA008884 | 1:1000 |
| Nfib | rabbit | Abcam | ab186738 | 1:100 |
| Onecut2 | sheep | R&D | AF6294 | 1:500 |
| Pou2f2 | rabbit | Abcam | ab178679 | 1:1000 |
| Sim1 | rabbit | Aviva SysBio | ARP33296_P050 | 1:50 |
| Sox2 | mouse | Santa Cruz Biotechnology | sc-365823 | 1:200 |
| Sox2 | goat | R&D | AF2018 | 1:500 |
| TH | sheep | R&D | AF7566 | 1:1000 |
| Zfhx3 | sheep | R&D | AF7384 | 1:1000 |
| Zfhx4 | rabbit | Atlas Antibodies | HPA023837 | 1:1000 |

**Table S2.** Related to Experimental Procedures: List of antibodies for flow cytometry.

| Antibody | Species | Company | Catalogue number | Dilution | secondary antibody |
| --- | --- | --- | --- | --- | --- |
| Neurod2 | rabbit | Abcam | ab104430 | 1:1000 | donkey-anti-rabbit A488 (Life Technologies) |
| Nfia | rabbit | Atlas Antibodies | HPA008884 | 1:1000 | donkey-anti-rabbit A488 (Life Technologies) |
| Nkx2.2-PE | mouse | BD Biosciences | 564729 | 1:50 | N/A |
| Onecut2 | sheep | R&D | AF6294 | 1:500 | donkey-anti-sheep A568 (Life Technologies) |
| Pax3-APC | mouse | R&D | IC2457A | 1:50 | N/A |
| Sox2-V450 | mouse | BD Biosciences | 561610 | 1:100 | N/A |
| Tubb3:A488 | mouse | BD Biosciences | 560381 | 1:100 | N/A |
| Tubb3:A647 | mouse | BD Biosciences | 560394 | 1:100 | N/A |
| Zxh3 | sheep | R&D | AF7384 | 1:500 | donkey-anti-sheep A568 (Life Technologies) |

**Table S3. Related to Figures 4,5,6,7: Summary of flow cytometry results** (Provided as separate Excel file)

**Table S4.**
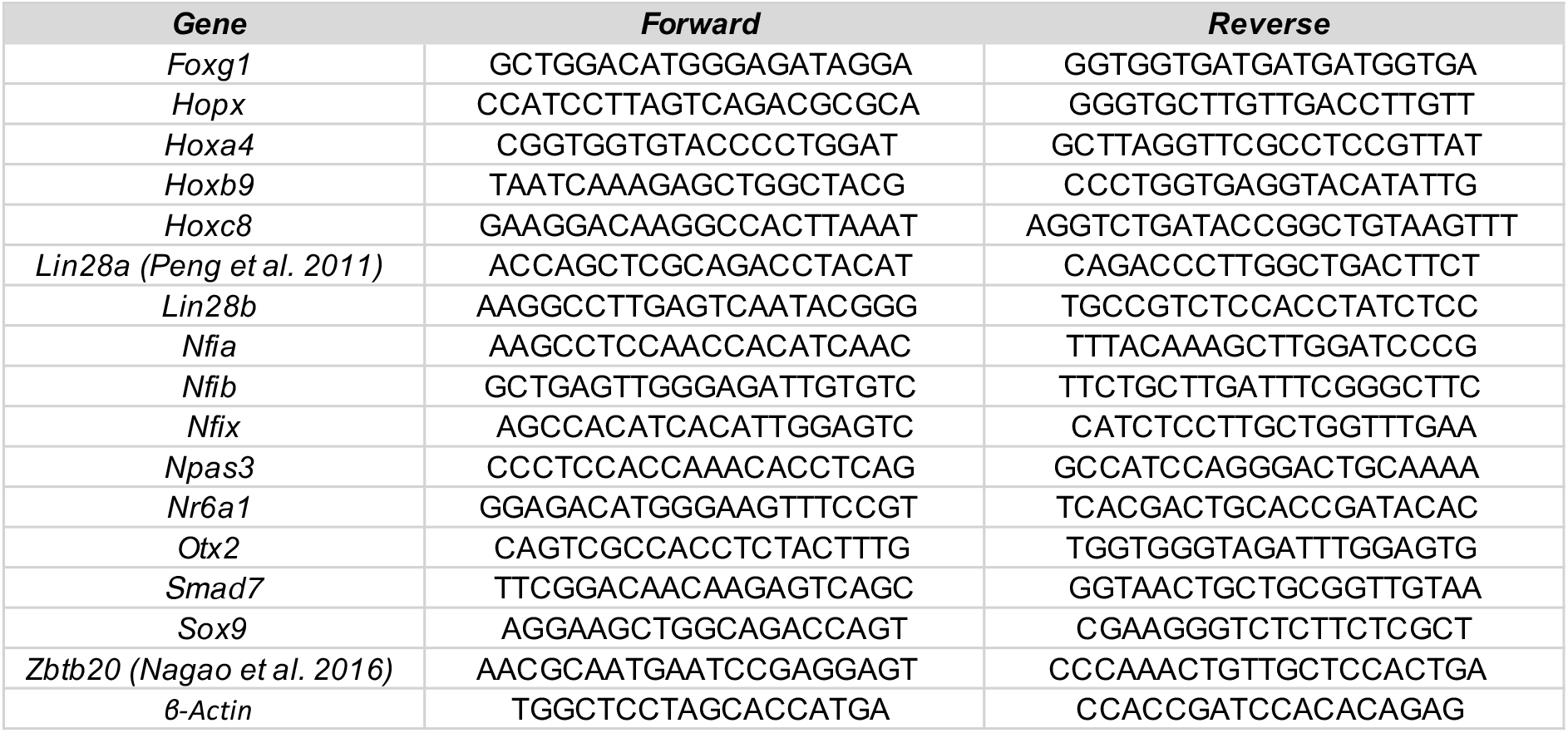
Related to Experimental Procedures: List of primers for RT-qPCR analysis.

